# Dually biofortified cisgenic tomatoes with increased flavonoids and branched-chain amino acids content

**DOI:** 10.1101/2022.10.31.514528

**Authors:** Marta Vazquez-Vilar, Asun Fernandez-del-Carmen, Victor Garcia-Carpintero, Margit Drapal, Silvia Presa, Dorotea Ricci, Gianfranco Diretto, José Luis Rambla, Rafael Fernandez-Muñoz, Ana Espinosa-Ruiz, Paul D. Fraser, Cathie Martin, Antonio Granell, Diego Orzaez

## Abstract

Higher dietary intakes of flavonoids may have a beneficial role in cardiovascular disease prevention. Additionally, supplementation of branched-chain amino acids (BCAAs) in vegan diets can reduce risks associated to their deficiency, particularly in older adults, which can cause loss of skeletal muscle strength and mass. Most plant-derived foods contain only small amounts of BCAAs and those plants with high levels of flavonoids are not eaten broadly. Here we describe the generation of metabolically-engineered cisgenic tomatoes enriched in both flavonoids and BCAAs. In this approach, coding and regulatory DNA elements, all derived from the tomato genome, were combined to obtain a herbicide-resistant version of an acetolactate synthase (mSlALS) gene expressed broadly, and a MYB12-like transcription factor (SlMYB12) expressed in a fruit-specific manner. The mSlALS played a dual role, as a selectable marker as well as being key enzyme in BCAA enrichment. The resulting cisgenic tomatoes were highly enriched in Leucine (21-fold compared to wild type levels), Valine (9-fold), Isoleucine (3-fold), and concomitantly biofortified in several antioxidant flavonoids including kaempferol (64-fold) and quercetin (45-fold). Comprehensive metabolomic and transcriptomic analysis of the biofortified cisgenic tomatoes revealed marked differences to wild type and could serve to evaluate the safety of these biofortified fruits for human consumption.

## Introduction

Personalized nutrition aims to improve life quality and prevent disease by customizing diets based on individual characteristics such as age, genetic background, gut flora composition, behavioral habits, etc. As our ability to generate individualized genetic and metabolomic profiles grows, so does the need to expand the assortment of reliable food sources from which important dietary compounds can be obtained. In recent times, the increasing precision offered by some new plant breeding techniques such as gene editing or cisgenesis, has contributed to raising public acceptance of plant biotechnology applied to food, as exemplified by the pioneering commercialization of GABA-enriched gene edited tomatoes (Waltz, 2021). At the same time, new breeding techniques open the way to the design of new metabolically customized fruits that combine non-competing metabolite enrichment strategies, narrowing the gap between precision nutrition and metabolic engineering. In this work, we describe the engineering of tomatoes dually enriched in flavonoids and branched-chain amino acids (BCAAs) using a cisgenic approach.

Flavonoids have a proposed role in the prevention of cardiovascular diseases and colon cancer (Martin et al., 2013). The overexpression of regulatory genes that activate several enzymes of the pathway was reported to lead to the accumulation at high levels of different flavonoid compounds such as anthocyanins, flavonols, or both, depending on the specific combination of transcription factors employed (Bovy et al., 2002; Butelli et al., 2008; Luo et al., 2008; Zhang et al., 2015). In particular, transgenic overexpression of Arabidopsis MYB12 (AtMYB12), a master regulator of the phenylpropanoid biosynthetic pathway, led to a substantial increase in flavonol levels in fruits (Luo et al., 2008).

There are increasing concerns about the effect that strictly vegetarian diets could have in the development of muscle related conditions, especially in older adults. Low Body Mass Index (BMI) and reduced muscle mass have been associated with the increased risk of hip fracture observed in women following vegetarian diets (Webster et al., 2022). Sarcopenia, a condition characterized by loss of muscle mass in older adults, is also a condition of concern associated with diets with low animal protein intake (Beaudart et al., 2017; Huang et al., 2016). BCAAs, valine (Val), leucine (Leu) and isoleucine (Ile), account for 14-18% of muscle protein total amino acids and they are considered critical regulators of anabolism in skeletal muscle tissues (Brestenský et al., 2015). Supplementation with these essential amino acids abundant in animal proteins is, alongside with resistance training, a standard treatment for sarcopenia (McKendry et al., 2020). Indeed, a lucrative market of BCAA supplements has grown in athletics due to the claims that relate these supplements with an increase in BMI (Kårlund et al., 2019). A more desirable solution to dietary supplements would consist in enriching fruits and vegetables in key compounds whose absence could bring long term negative effects in vegetarian diets. In this regard, elevating BCAA content in fruit and vegetables could reduce problems associated with the highly recommended reduction in the intake of animal-based food in older adults, especially for individuals opting for a strict vegan diet (Domić et al., 2022; Reid-McCann et al., 2022). An unexplored strategy to increase BCAAs in plant tissues consists of overexpression of the acetolactate synthase (ALS), an enzyme that catalyzes the first step of BCAA biosynthesis, in particular the conversion of pyruvate into 2-acetolactate to Val and Leu and the pyruvate conversion into 2-aceto-2-hydroxybutyrate, a precursor of Ile.

Cisgenesis is a new plant breeding technique (NPBT), that stands astride classical breeding and transgenesis (Espinoza et al., 2013). Cisgenesis makes only use of the genetic pool from a plant species itself or cross-compatible ones for engineering crops with new agronomic traits. By avoiding the introduction of alien DNA in the crop’s genome, cisgenic approaches aim to reduce the biosafety concerns associated to transgenesis. Cisgenesis requires the use of compatible selection procedures. Some selectable markers have been derived from plants and those conferring resistance to herbicides can be adapted to cisgenic approaches (Liu et al., 2013; Sundar & Sakthivel, 2008; Tian et al., 2015; Yu et al., 2015), including the use of mutated versions of the above mentioned ALS enzyme (Okuzaki et al., 2007; Shimizu et al., 2008; Yao et al., 2013). ALS inhibition by herbicides results in a reduction of BCAAs in the plant and ultimately in plant death (Binder, 2010). Different amino acid substitutions have been reported to confer tolerance to different ALS-inhibiting herbicides by blocking the entrance of pyruvate to the active site of the enzyme (Zhou et al., 2007). One of them, the Pro-197-Ser mutation (in the ALS of *Arabidopsis thaliana*) results in sulphonylurea resistance (Haughn et al., 1988). Selection markers based on this mutated ALS have been developed for crops such as apple (Yao et al., 2013) and tobacco (Haughn et al., 1988), but an equivalent strategy remains unavailable for tomato.

We show here that the generation of tomatoes dually-biofortified in flavonoids and BCAAs can be achieved by combining the constitutive expression of a mutated version of a *Solanum lycopersicum* ALS (mSlALS) gene and the previously described tomato MYB12 gene (SlMyb12) (Ballester et al., 2010) driven by a fruit-specific promoter. In this approach, ALS played a dual role, both as cisgenic selectable marker and as key factor for BCAA enrichment. Metabolic analyses demonstrate that cisgenic fruits were enriched in flavonols with significant accumulation of rutin, quercetin and kaempferol in the fruit flesh. Additionally, our data show that overexpression of mSlALS results, not only on herbicide resistance but also in increased content of BCAAs and BCA-devived volatiles. Finally, we used transcriptomics and metabolomics to provide an unbiased picture of the effects in the fruit of the simultaneous expression of SlMYB12 and SlALS.

## Results

### Design of a new strategy for tomato cisgenic biofortification

An all-tomato-based selectable marker requires an endogenous gene expressed under tomato regulatory sequences. We aimed to design a cisgenic selectable marker based on ALS. A BLAST search with the AtALS amino acid sequence resulted in the identification of the three putative ALS homologs in tomato previously reported by Gao et al. (2014), ALS1 (Solyc03g044330.1), ALS2 (Solyc07g061940.2) and ALS3 (Solyc06g059889.2). Two of them, ALS1 and ALS2, have the highest amino acid sequence similarity with AtALS. We selected ALS1 to set up a cisgenic selection marker for tomato transformation. A single nucleotide change (C to T at position 556), resulting in a proline to serine mutation at amino acid 186 (position 197 in reference to AtALS) was introduced to create mSlALS (Figure 1A). In parallel, tomato promoter regions conferring high and constitutive expression were isolated from genes showing high and broadly distributed expression in the Tomato Functional Genomics Database (http://ted.bti.cornell.edu/). Three top candidate genes (Solyc01g099770, Solyc06g007510 and Solyc09g010800) were selected, and their putative 5’ regulatory regions (comprising the annotated 5’ UTRs plus a 2 kb fragment upstream of the transcriptional start site (TSS), and 3’ regulatory regions (comprising the 3’ UTRs plus 1 kb fragment downstream the transcriptional termination site) were isolated and tested transiently in *Nicotiana benthamiana* using a Firefly luciferase (FLuc) assay as previously described by Vazquez-Vilar et al. (2017). Among the different assayed promoters, the metallothionein-like protein type 2B (Mtb, Solyc09g010800) promoter, in combination with its own terminator, was shown to confer the highest relative expression levels, 6.94 ± 0.95 RPUs (Figure 1B, Sarrion-Perdigones et al., 2013). The transcriptional activity conferred by the Mtb promoter and terminator regions in the assayed conditions was approximately half of the activity conferred by the strong CaMV35S promoter when combined with the nopaline synthase (nos) terminator. Based on these data, we assembled the mSlALS CDS with the Solyc09g010800 promoter and terminator regions generating the transcriptional unit (TU) pMtb::mSlALS::tMtb, ready to be used as a cisgenic selectable marker.

**Figure 1.**
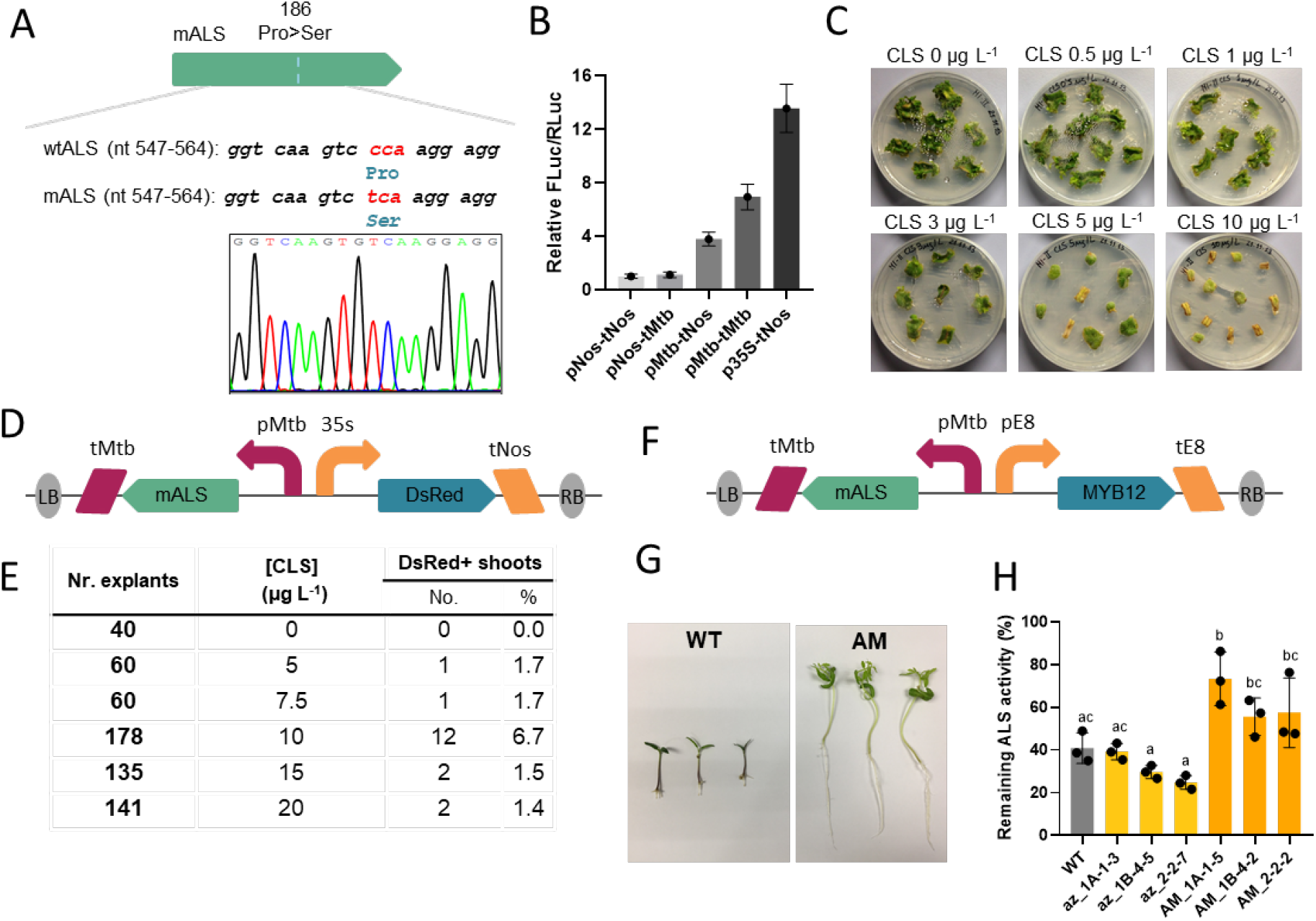
Overexpression of a mutated version of the acetolactate synthase gene confers chlorsulfuron resistance in tomato. A) Schematic representation and confirmatory chromatogram of the nucleotidic mutation introduced in the ALS that results on a Pro>Ser amino acid change at position 186. B) Relative Firefly Luciferase (FLuc) / Renilla Luciferase (RLuc) activity conferred by different promoter and terminator combinations (Sarrion-Perdigones et al., 2013). C) Untransformed cotyledon explants maintained for 5 weeks on regeneration medium supplemented with different chlorsulfuron concentrations. D) Genetic construct used for optimization of the tomato transformation protocol with the mALS selection marker. E) Number of transgenic shoots recovered with different CLS concentrations in a transformation carried out with the genetic construct depicted in D). F) Schematic representation of the genetic construct used for the generation of the ALS-MYB (AM) lines. G) 10-days old seedlings of WT and cisgenic AM lines grown in MS supplemented with 50 µg/L chlorsulfuron. H) Calculated remaining ALS activity in leaf extracts of three independent T2 AM lines and their corresponding azygous lines in the absence and presence of 50 µg/L chlorsulfuron. Error bars represent SD of independent biological replicates (n=3). Statistical analyses were performed using one-way ANOVA (Tukey’s multiple comparisons test, P-Value ≤ 0.05). Variables within the same statistical groups are marked with the same letters.

Next, we established a protocol for tomato transformation with the new marker. First, we tested shoot regeneration of untransformed *S. lycopersicum* cv. Moneymaker cotyledon explants at different chlorsulfuron (CLS) concentrations (Figure 1C). Based on these results, we carried out the optimization of the new cisgenic marker for Agrobacterium-mediated transformation using a linked DsRed fluorescent reporter construct (Figure 1D), and testing a range of CLS concentrations in the regeneration media. The number of transformed explants was calculated based on their DsRed fluorescence. The optimal CLS concentration was set at 10 µg L^−1^ resulting in a transformation efficiency of 6.7% (Figure 1E).

Finally, to obtain mature fruits cisgenically biofortified in flavonoids, we produced a construct where the expression of an endogenous MYB12 homologue (SlMYB12, coding sequence of Solyc01g079620.3) was driven by the ethylene-responsive, fruit-specific regulatory regions of the tomato E8 gene (Solyc09g089580.3.1). The assembly of this new TU next to the cisgenic ALS marker resulted in the genetic module tMtb::mSlALS::pMtb-pE8::SlMYB12::tE8 (Figure 1F). Eleven primary transformants were obtained after transformation with this genetic construct from 200 explants (5.5% transformation efficiency). T1 seeds of three of the lines, hereafter referred to as AM lines, were selected in MS medium supplemented with 10 µg L^−1^ CLS. Only plants carrying the T-DNA grew in these conditions, confirming the stability of the cisgenic marker (Figure 1G). All three AM lines showed a segregation corresponding to the insertion of a single copy of the T-DNA (data not shown). To further corroborate the CLS resistance observed in seedlings, we determined the ALS enzymatic activity in leaves of AM and wild type plants. As shown in Figure S1, all AM, azygous and WT plants showed detectable ALS activity, but only AM samples partially retained ALS activity when incubated in the presence of CLS (Figure 1H). The partial loss of ALS activity in cisgenic plants in the presence of the herbicide can be explained by the presence of two additional wild type ALS gene copies in the genome which are susceptible to CLS. Six T1 plants from each AM line were grown to maturity and self-pollinated to the next generation. All experiments described below were carried out in plants of the T2 and subsequent generations.

### AM cisgenic tomatoes show an enrichment of BCAA metabolism in fruit

The expression levels of both cisgenic (cALS) and endogenous (eALS) transcripts were confirmed using a RT-qPCR analysis with primers specifically designed to differentiate between the two forms. cALS transcript showed strong overaccumulation in fruit. Furthermore, cALS mRNA levels showed an unexpected 25-fold increment from green to orange stages (Figure 2A). These data led us to investigate the extent to which the high levels of ALS transcript in the fruit flesh would affect the fruit composition, and in particular, the amount of BCAAs and their derivatives. We found much higher amounts of leucine, isoleucine and valine in the three lines under analysis (Figure 2B). For AM_2-2-2, increments were 14, 3.8 and 7.8-fold for Leu, Ile and Val, respectively, with respect to WT fruits. Absolute contents of all three BCAAs in fruits of line AM_2-2-2 are shown in Table 1. As can be observed, the accumulation of free leucine reached up to 350 mg/kg FW (estimated 7.0 g/kg dry weight), whereas more modest accumulations of isoleucine (45 mg/kg FW, estimated 0.9 g/kg DW) and valine (140 mg/kg FW, estimated 2.8 g/kg DW) were measured. To discard a possible effect of MYB12 in the changes observed in BCAAs in the fruit, we studied levels of Leu, Ile and Val also in the leaves of the three lines. As expected, BCAAs were found to overaccumulate also in the leaves of cisgenic plants. AM_1A-1-5 and AM_2-2-2 leaves showed the highest increments in BCAAs (Figure S2). For AM_1A-1-5, increments were 2.7, 4.7 and 6.7-fold WT for Leu, Ile and Val, respectively.

**Figure 2.**
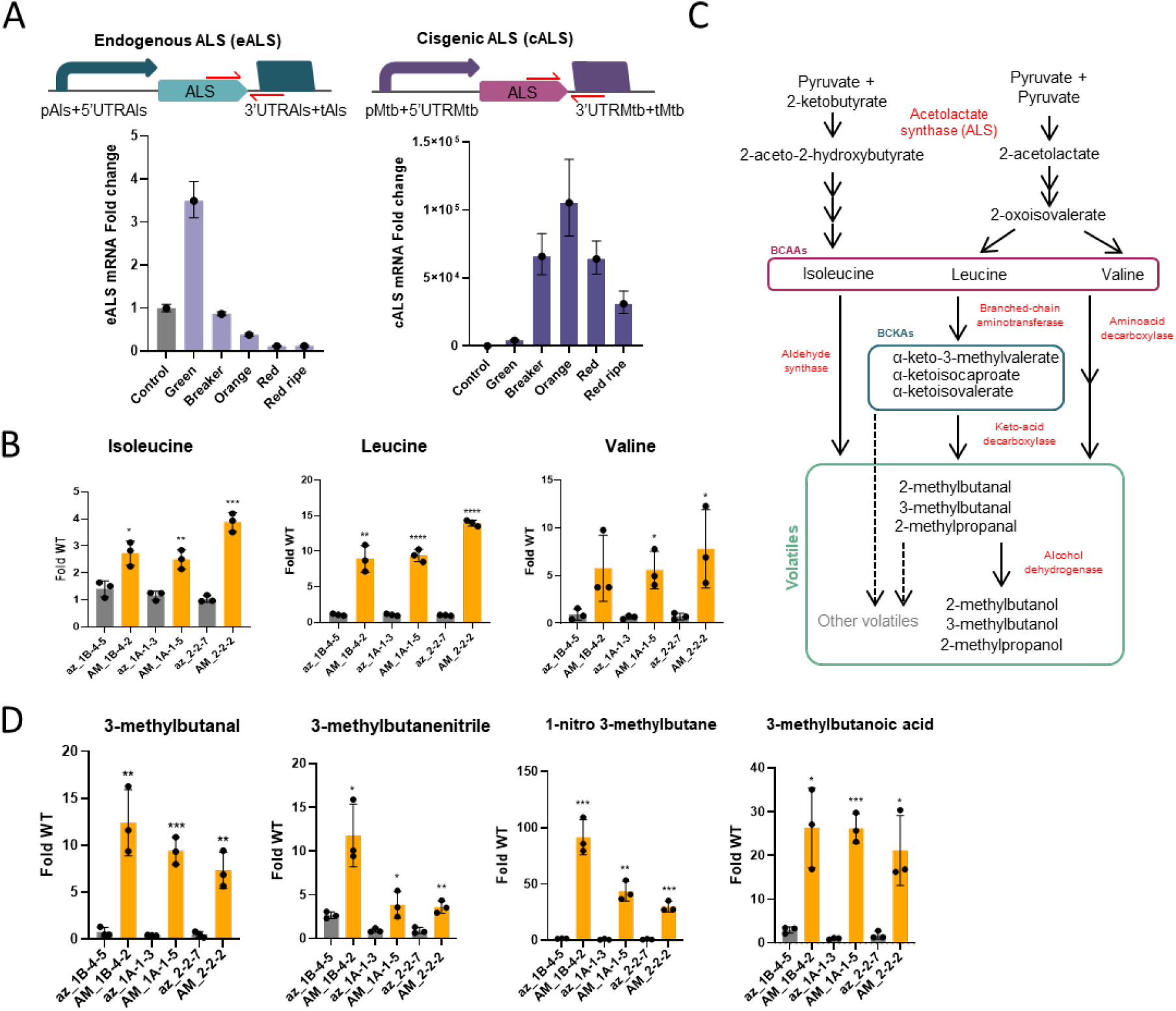
Overexpression of the acetolactate synthase gene results in increased BCAAs and branched chain volatiles in tomato fruits. A) Primer design for differential detection of the endogenous and the cisgenic *Solanum lycopersicum* ALS mRNA (top) and mRNA fold change of endogenous and cisgenic ALS (bottom). Fruits of a T3 2-2-2 homozygous plant were harvested at different ripening stages. The control sample is a fruit of an azygous line harvested at breaker. Error bars represent standard deviation of technical replicates (n=3). B) Schematic representation of the BCAAs biosynthetic pathway and proposed pathways for branched chain volatiles synthesis. C) Fold change on the BCAAs content of red ripe tomato fruits of three independent T2 AM lines compared to their corresponding azygous lines. D) Fold change on the branched chain volatiles content of red ripe tomato fruits of three independent T2 AM lines compared to their corresponding azygous lines determined using GC-MS. Asterisks indicate values that are significantly different *(P<0.05), **(P<0.01), ***(P<0.001) from their corresponding azygous lines (Student’s t-test). Error bars represent SD of independent biological replicates (n=3).

**Table 1.** Absolute quantification of BCAAs in flesh of three independent AM and three independent WT fruits collected at 7dpb.

|  | WT-1 | WT-2 | WT-3 | AM-1 | AM-2 | AM-3 |
| --- | --- | --- | --- | --- | --- | --- |
| **Isoleucine**<br>**($\mu\text{g/g FW}$)** | 16.23 ± 1.43 | 16.26 ± 1.73 | 10.63 ± 0.30 | 42.76 ± 2.70 | 49.38 ± 1.75 | 43.54 ± 1.72 |
| **Leucine**<br>**($\mu\text{g/g FW}$)** | 17.73 ± 1.70 | 17.48 ± 1.73 | 11.29 ± 0.20 | 299.78 ± 17.68 | 358.26 ± 18.60 | 306.58 ± 10.22 |
| **Valine**<br>**($\mu\text{g/g FW}$)** | 14.82 ± 0.91 | 14.61 ± 2.12 | 10.57 ± 0.47 | 99.85 ± 21.12 | 137.04 ± 2.11 | 120.80 ± 3.43 |

Branched chain volatiles (BCVs) share with BCAAs the same biosynthesis metabolic pathway (Kochevenko et al., 2012; Rambla et al., 2014). This group of volatiles is highly relevant for tomato fruit liking as, of the circa 30 volatiles considered to be involved in the human perception of tomato flavor, 8 of them are BCVs: 3-methylbutanenitrile, 1-nitro-3-methylbutane, 3-methylbutanoic acid, 3-methylbutanol, 3-methylbutanal, 2-isobutylthiazole, 2-methylbutanol and 2-methylpropyl acetate (Tieman et al., 2017). Twelve BCVs were identified in the fruit via GC-MS. Four of them, 3-methylbutanal, 3-methylbutanenitrile, 3-methylbutanoic acid and 1-nitro-3-methylbutane, were significantly increased in the three cisgenic lines. The biosynthesis of these four volatiles is related to the BCAA leucine, which was also the amino acid with the most altered content. The most dramatically altered BCVs were 1-nitro-3-methylbutane and 3-methylbutanoic acid, respectively showing up to 91.5 and 26.3-fold increments as compared to the WT (Figure 2D). These increases are remarkably higher than those observed in the BCAAs, thus suggesting a looser control in the accumulation of these leucine-related volatiles as compared to their respective amino acid. Other BCVs, including all those related to isoleucine and valine, showed either significant increase only in some of the lines or non-significantly altered levels (Figure S3).

### Fruit-specific expression of SlMYB12 enhances flavonoids production

The characterization of the cisgenic lines proceeded with the validation of the functionality of the second TU present in the AM lines (pE8::SlMYB12::tE8). We first verified the E8-mediated ripening-dependent expression of the cisgenic SlMYB12 gene (cMYB12), using RT-qPCR primers pairs that discriminate between cisgenic and endogenous (eMYB12) copies. As expected, cMYB12 was almost undetectable in green fruits and its levels sharply increased at breaker (220-fold increment), reaching a maximum at orange stage, and decreasing at later maturation stages (Figure 3A). In comparison, eMYB12 expression showed smaller variations with ripening (13-fold increase from green to breaker), following a decreasing trend at later stages (Figure 3A). While cMYB12 could be detected both in the flesh and in the peel of cisgenic fruit, eMYB12 was only expressed in the peel, both in WT and AM fruits (Figure S4).

**Figure 3.**
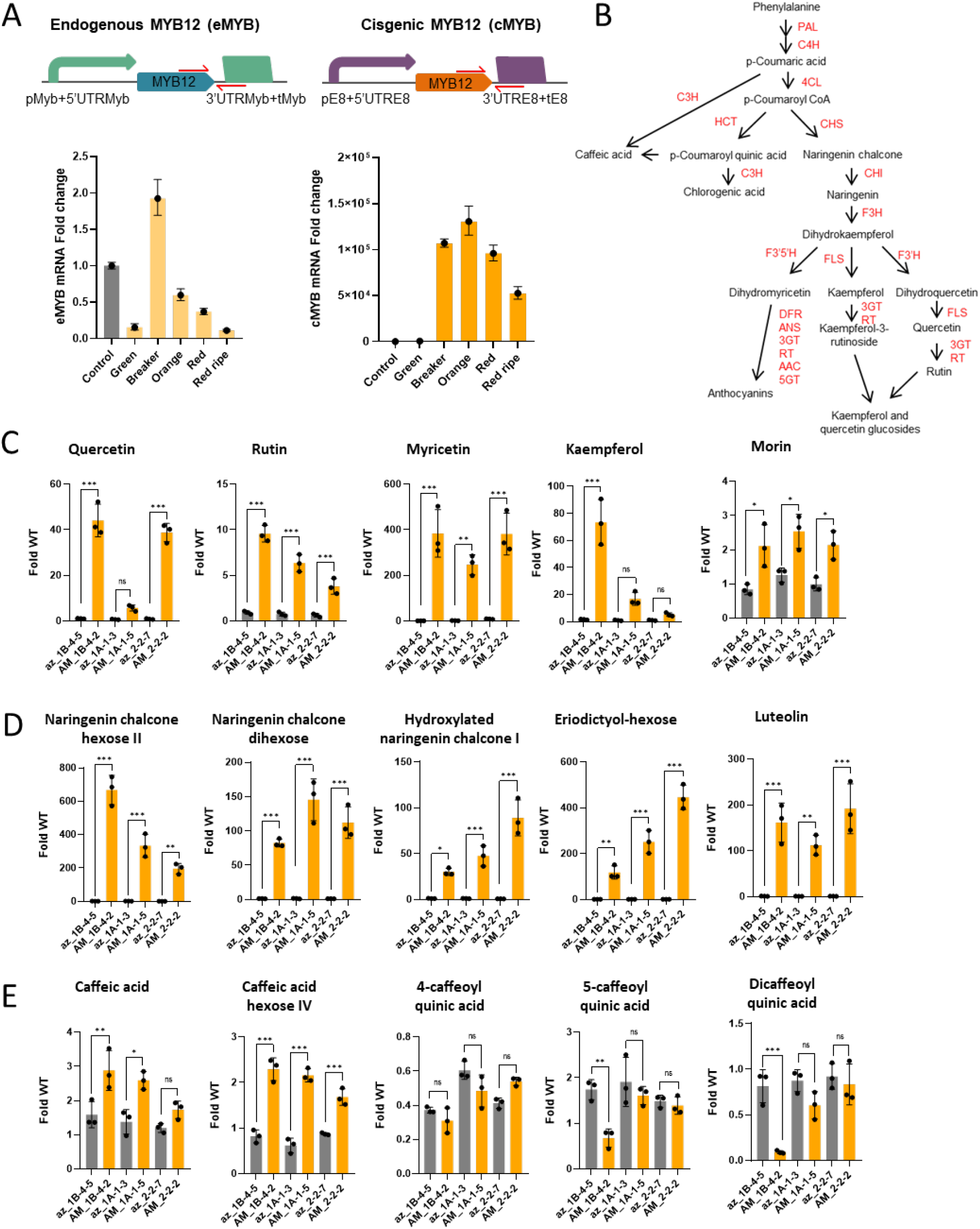
Fruit-specific expression of SlMYB12 results in increased phenylpropanoids content. A) Primer design for differential detection of the endogenous and the cisgenic Solanum lycopersicum ALS mRNA (top) and mRNA fold change of endogenous and cisgenic ALS (bottom). Fruits of a T3 2-2-2 homozygous plant were harvested at different ripening stages. The control sample is a fruit of an azygous line harvested at breaker. Error bars represent standard deviation of technical replicates (n=3). B) Schematic representation of the phenylpropanoids biosynthetic pathway. C) Fold change on the flavonols content of red ripe tomato fruits of three independent T2 AM lines compared to their corresponding azygous lines determined using HPLC. D) Fold change on other flavonoids content of red ripe tomato fruits of three independent T2 AM lines compared to their corresponding azygous lines determined using HPLC. E) Fold change on the caffeoyl quinic acids content of red ripe tomato fruits of three independent T2 AM lines compared to their corresponding azygous lines determined using HPLC. Asterisks indicate values that are significantly different *(P<0.05), **(P<0.01), ***(P<0.001) from their corresponding azygous lines (Student’s t-test). Error bars represent SD of independent biological replicates (n=3).

As a final characterization step, we analyzed the content of several compounds in the flavonoid pathway (Figure 3B) in AM and control fruits. For this, red ripe fruits from three different cisgenic plant lines (1B-4-2, 1A-1-5, and 2-2-2), together with fruits of their corresponding azygous lines (1B-4-5, 1A-1-3, and 2-2-7, respectively), and a control wild type plant were separately analyzed by LC-ESI (+/-)-MS, and the most relevant flavonoids were identified and quantified relative to the WT. Major differences were observed in several specific flavonol compounds in line AM_1B-4-2, where quercetin, rutin, myricetin, morin and kaempferol showed over-accumulation levels of 40-, 10-, 400-, 2- and 70-fold respectively (Figure 3C). Significant over-accumulations were found also for the glycoside derivatives of these compounds, including kaempferol-diglucoside, kaempferol-dihexose, kaempferol-dihexose deoxyhexose, kaempferol-glucose rhamnose, kaempferol-glucosyl-glucoside-rhamnoside, kaempferol-rutinoside, quercetin hexose and quercetin deoxyhexose-hexose-deoxyhexose (Figure S4). In addition to flavonols, other flavonoid compounds including flavanones such as naringenin chalcone hexose, naringenin chalcone dihexose, naringenin chalcone glucoside, hydroxylated naringenin chalcone, eriodictyol-hexose and methyl-ether of (eriodictyol /eriodictyol chalcone) hexose, and flavones such as luteolin were also over-accumulated in AM fruits (Figure 3D, Figure S5).

As Luo et al. reported modifications on caffeoyl-quinic acids (CQAs) levels when AtMYB12 was expressed in tomato fruits (Luo et al., 2008), we decided to investigate whether these compounds were also modified in the AM lines. We observed a modest 2.9- and 2.6-fold enrichment of caffeic acid for lines 1B-4-2 and 1A-1-3, respectively, while for line 2-2-2 no substantial modification on caffeic acid levels was detected. For all three lines, an increment in caffeic acid hexose IV of roughly 2-fold was observed. However, no differences were observed in any of the lines for the CQAs content. Finally, we carried out absolute quantification of most relevant flavonol aglycones quercetin and kaempferol. As shown in Table 2, the enrichment of fruits in this type of compound was confirmed in absolute values quantifications, with remarkable accumulations that reach up to 100 µg / g FW of quercetin in ripe fruits.

**Table 2.** Absolute quantification of flavonols in flesh of three independent AM and three independent WT fruits collected at 7dpb.

|  | WT-1 | WT-2 | WT-3 | AM-1 | AM-2 | AM-3 |
| --- | --- | --- | --- | --- | --- | --- |
| **Quercetin**<br>**($\mu\text{g/g FW}$)** | 1.73 ± 0.03 | 2.00 ± 0.07 | 1.84 ± 0.05 | 97.53 ± 3.16 | 80.96 ± 4.52 | 74.85 ± 2.27 |
| **Kaempferol**<br>**($\mu\text{g/g FW}$)** | 0.216 ± 0.004 | 0.259 ± 0.001 | 0.242 ± 0.019 | 14.43 ± 0.67 | 16.65 ± 0.60 | 14.70 ± 1.55 |

### Metabolic fingerprinting clearly separate AM tomatoes from WT both in peel and flesh

A general view on how this new metabolic engineering approach affected the overall fruit metabolite composition was obtained by means of an untargeted LC-MS metabolomic profiling. Principal component analysis (PCA) of metabolic features detected in flesh and peel samples collected at 4 and 7 dpb showed a clear discrimination between AM and WT samples (Figure 4A). Class separation becomes more evident at the later ripening stages both in peel and flesh, as expected for the ripening-associated expression programmed for Myb12. A heatmap of the 100 most statistically different features showed that the most relevant changes observed in fruits corresponded to metabolites whose relative abundance increases in AM fruit, while only a few features increased their relative accumulation levels in the WT (Figure 4B). This bias towards metabolite accumulation occurs in flesh samples both at 4 dpb and at 7 dpb. In contrast, the direction of changes seemed more balanced in peel, with even a majority of downregulated features observed at 7 dpb. A Venn diagram of features that were differentially overexpressed in AM shows that approximately 80% of the differential features were present in the flesh and a 68% of them in the peel (Figure 4C). Among the features overexpressed in WT, the 83% of them are present in the peel and the 66% in the flesh. 136 (30%) were present in flesh and peel collected at 7 dpb and 108 (24%) were present only in the peel at 7 dpb.

**Figure 4.**
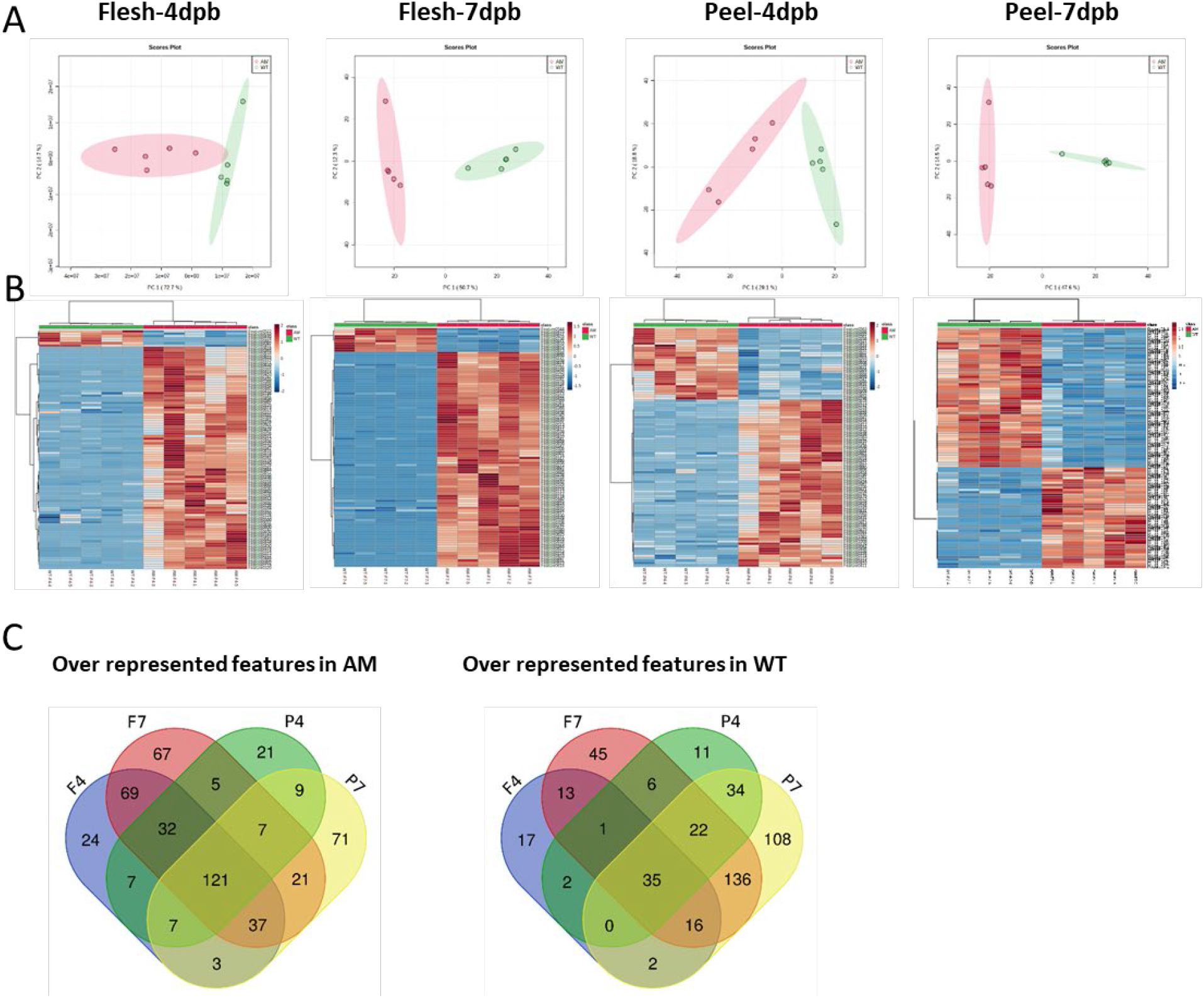
Metabolite profiling with LC-MS of ALS-MYB flesh and peel samples. A) Principal component analysis (PCA) of AM versus WT samples. B) Heatmap of the 100 most differential features detected in AM and WT flesh and peel samples. C) Venn diagram with the number of independent and common features among the different samples.

#### Transcriptomic analysis of AM fruits revealed a general activation of flavonoids and branched amino-acid pathways

Further investigation of the effects of simultaneous expression of ALS and MYB at high levels in the fruit was carried out using transcriptomic analysis. Tomato fruits of AM and WT plant lines were harvested at 4 dpb and flesh samples were used for RNAseq analysis. In the flesh, a total of 1874 genes were differentially expressed, a majority of then (1200) overexpressed in the AM fruits (Table S1). Interestingly, SlALS ranked second among the most significant upregulated genes. We estimated that in the cisgenic line ∼99.2% of reads corresponded to the cisgenic copy and only ∼0.8% of them to the endogenous one. GO term enrichment analysis revealed that genes involved in the small molecule, carboxylic acid, oxoacid, organic acid, nucleoside/nucleotide and cellular amino acid biosynthesis processes were significantly overrepresented in the AM fruits (Figure 5A and Table S2). Regarding the cellular component, the GO terms significantly overrepresented in the AM lines were mainly related to the plastids and their different compartments (Figure 5A and Table S2). The list of genes contributing to these GO terms include the acetolactate synthase and several plastidial kinases and oxidoreductases involved in the shikimate and phenylpropanoids pathways (see Table S3).

**Figure 5.**
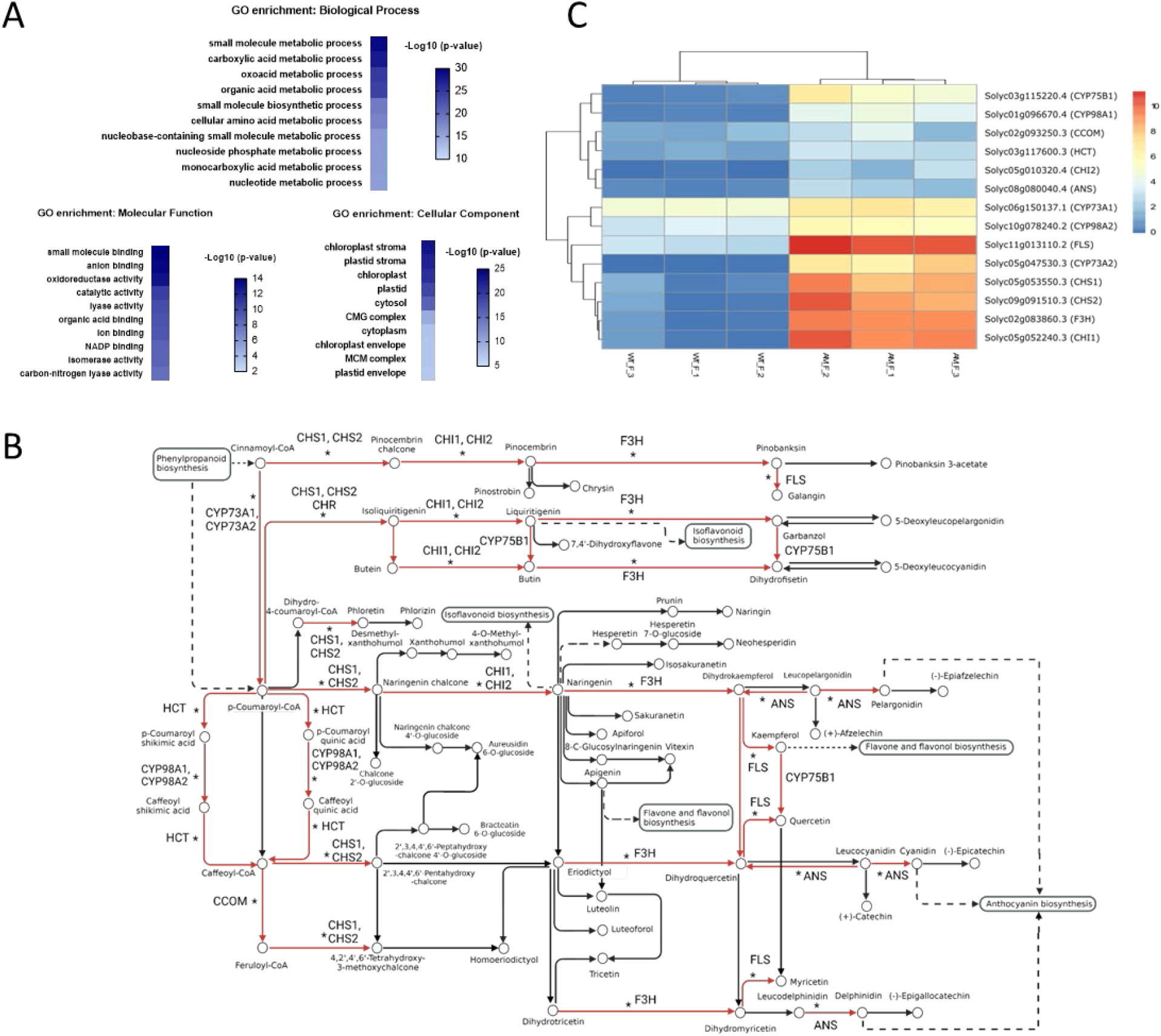
Transcriptomic analysis of ALS-MYB flesh samples shows overexpression of multiple genes involved in the amino acids and flavonoids biosynthetic pathways. A) GO terms enrichment analysis of genes overexpressed in the AM fruits. The ten most significant GO terms are displayed. B) A KEGG analysis shows that several genes of the flavonoids pathway (red arrows) are overexpressed in the AM fruits compared to WT. Asterisks indicate genes overexpressed in AtMYB12 fruits from Zhang et al. (2015). C) Log FPKMs of the genes differentially expressed among lines in the flavonoids pathway depicted in B).

A KEGG enrichment analysis confirmed the biosynthesis of secondary metabolites as the most significantly enriched biological process, next to the amino acids biosynthesis. A closer look to the flavonoid pathway revealed a total of 14 upregulated genes in AM fruits, distributed in multiple steps of the pathway (see Figure 5B). Among them, the strongest upregulation occurred in chalcone synthases 1 and 2 (CHS1 and CHS2), the naringenin 3-dioxygenase (F3H), the chalcone isomerase 1 (CHI1) and the flavonol synthase (FLS) (Figure 5C). This latter gene, catalyzing the oxidation of dihydroflavonols to produce flavonols, showed the most dramatic upregulation, in accordance with the observed increase of kaempferol, quercetin and myricetin. The KEGG analysis also revealed that 50 genes involved in the amino acid biosynthesis pathway were differentially expressed in the AM fruits, most of them upregulated (Figure S6). Interestingly, those upregulated genes in the shikimate pathway leading to the biosynthesis of phenylalanine were also identified by Zhang et al. (2015) upon ectopic expression of AtMyb12 driven by E8 promoter, whereas others leading to BCAAs synthesis are exclusive of the combination of SlMyb12 and SlALS.

## Discussion

Cisgenesis refers to those genetic engineering approaches that make use only of DNA elements derived from the genetic pool of sexual compatible species. Cisgenic crops are intrinsically genetically modified organisms (GMOs), and therefore they are sensibly regulated as such (van Hove & Gillund, 2017). However, bearing in mind the reduction in perceived risk associated with the avoidance of exogenous DNA sequences in the makeup of cisgenic plants by consumers, it is reasonable to claim and to expect a reduction in the regulatory constraints governing their consumption and/or commercialization, especially in those countries where process-based GMO regulations prevail (Russell & Sparrow, 2008). Interestingly, several surveys show that public acceptance is higher for crops generated using new breeding technologies that avoid the presence of foreign DNA sequences (Delwaide et al., 2015). Recently, the EU Commission conducted a public have-your-say enquiry on the need to change regulations for plants produced using new breeding techniques, a survey that was focused on genome editing but that extended to cisgenesis. The expectation for regulatory changes has brought along a renewed interest in the evaluation of cisgenesis-based engineered crops. An immediate application of cisgenesis is the enrichment of food composition in health-related compounds, more specifically in those compounds that are already present in the crop, but for which genetic intervention could result in a higher abundance or a more convenient relative concentration. The cisgenic substitution of the regulatory sequences governing the expression of transcription factors (TFs), or biosynthetic enzymes (BEs) are two strategies to modify metabolic fluxes leading to the accumulation of health-promoting compounds. Here, we combined the TF and BE approaches in a single cisgenic intervention in tomato fruits, demonstrating the feasibility of combined fortification strategies.

The herbicide-resistance trait conferred by mutations in ALS loci has been used traditionally in many crops (Okuzaki et al., 2007; Shimizu et al., 2008), with resistant mutations known to appear spontaneously in weed species (Tranel & Wright, 2002). The employ of ALS as transgenic selection marker is also common (Z. Li et al., 1992; Yao et al., 2013), but its use in cisgenic breeding requires species-specific adaptation, limiting its widespread use for this particular purpose. Here, we showed that the 186 Pro>Ala conversion (197 in reference to AtALS) of SlALS confers resistance to chlorsulfuron, as previously demonstrated for the Arabidopsis ALS in tobacco (Haughn et al., 1988) and with the apple ALS both in tobacco and apple (Yao et al., 2013). The transformation efficiencies of 5.5% reported here were lower than those obtained by the use of the well-established nptII antibiotic selection marker but were high enough to be considered for routine transformations, particularly if the cisgenic status conferred a commercial advantage to the resulting product. To our knowledge no other ALS-based purely cisgenic breeding strategy has been described so far. Surprisingly, an associated increase in BCAAs has not been reported before, despite the extended use of this marker in transgenic selection strategies. A possible explanation is that the levels of expression of the mutated ALS in target food tissues were considerably lower to those conferred by the MTB promoter employed here. In other examples such as in apple (Yao et al., 2013), where ALS expression was driven by a CaMV35S promoter, the effect on BCAAs might have remained unnoticed. When the goal is to obtain the double biofortification, the choice of a strong promoter seems the best way to proceed, however in the light of our findings the employment of ALS under a strong promoter should be reconsidered when the aim is to generate new plants varieties free of effects related to the use of the selectable marker. For these cases, the use of endogenous ALS promoter could be an advisable strategy. Conversely, ALS-based BCAA enrichment could be also considered as an interesting biofortification strategy to apply to protein-rich crops to modify the profile of amino-acids in plant-based foods. Our results also suggest that transaminases and other enzymes involved in the BCV synthesis are not limiting factors, and by simply increasing the pool of BCAA precursors, higher levels of BCVs can be obtained. This point is reinforced by data generated by Breitel et al., (2020), which shows that E8-driven expression of AtMYB12 alone had no contribution to the BCAA increase in the fruit. Furthermore, we also observed strong accumulation in BCAAs-derived volatiles in leaves of cisgenic tomato lines. Since E8-driven gene expression is strictly fruit-specific, these observations strongly suggest that the observed BCAAs accumulation is solely due to the effect of ALS overexpression.

As expected, the parallel overexpression of SlMYB12 driven by E8 promoter resulted in a notable over-accumulation of specific compounds of the flavonol biosynthesis pathway in the fruit, in line with previous results reported with transgenic lines overexpressing the orthologous transcription factor from Arabidopsis (Luo et al., 2008). All three independent lines analyzed here showed a drastic increase of naringenin, kaempferol and quercetin glycosides in the fruit in reference to wild type and the corresponding azygous lines. The degree of enrichment in target metabolites obtained with cisgenesis seems comparable than that reported in equivalent transgenesis, despite the use of endogenous TF and the concomitant enrichment in BCAAs. Overaccumulations of most relevant aglycon flavonols such as quercetin and kaempferol, are close to 50-fold as compared with wild type Moneymaker fruits, a similar range to what was found earlier in transgenic tomatoes transformed with E8-driven Arabidopsis AtMYB12 gene (Zhang et al 2015), who established earlier the role of MYB12. Also in this previous study, Zhang et al. (2015) showed that AtMYB12 under the control of the E8 promoter resulted in transcriptomic changes in nearly all the genes involved in glycolysis, the pentose phosphate, the shikimate and the flavonoid pathways. Our data indicates that SlMYB12 is functionally equivalent to AtMYB12. We observed that, like AtMYB12, SlMYB12 has a role in the transcriptional activation not only of phenylpropanoid biosynthetic enzymes but also in rewiring carbon flux towards the production of aromatic amino acids. As expected, a correlation between up-regulated transcripts and metabolite levels was found for most of the flavonoid pathway steps when data were analyzed in the context of metabolic pathways.

In addition to the expected enrichment in targeted compounds, untargeted metabolomic profiles showed a clear separation between cisgenic and control fruits. Although the most distinctive features in untargeted profiles are likely to correspond to flavonols and BCAAs, both metabolomic analysis and comparative transcriptomics suggested that compositional changes go beyond these two main metabolic pathways. Separation among samples is more evident at the later ripening stage (7 dpb), arguably due to the late expression of the two cisgenes driven by the E8 and MTB promoters respectively. The untargeted metabolome profiles shown here highlight the extent of the compositional modifications that can be obtained by simply introducing changes in the expression levels of two unique genes, and this is expected to happen regardless of the use of transgenesis, cisgenesis or traditional breeding. In our view, this reflects the level of interconnection that occurs among biosynthetic pathways and questions the applicability of the substantial equivalence concept (Catchpole et al., 2005) for the evaluation of safety profiles in plant breeding when nutritional enrichment is the objective. It would be more informative to compare the metabolomic profiles of the engineered crop with other varieties of the same or related crops (e.g., tomatoes in this case), an approach that could help to identify any suspicious, unintended deviation from what is generally considered as safe in the metabolic composition of the same range of product. In this regard, the availability of increasingly complete pan-metabolomes from different crop species is a highly valuable resource (Drapal et al., 2022; Enfissi et al., 2021). In our view, strong unintended deviations are especially unlikely to occur using cisgenic breeding approaches.

Although dually fortified in flavonols and BCAAs, the metabolite enrichment levels in terms of overall dietary requirements are not comparable for the two types of metabolites. The absolute levels of quercetin produced in cisgenic fruits reached remarkable levels of 100 µg/g FW, equivalent to 10 mg of quercetin in an average 100 g serving portion, a quantity close to that employed when quercetin is used as a food supplement. On the other hand, the content of BCAAs in a serving portion of tomato represents no more than 2-5% of daily dietary requirements (Kurpad et al., 2006). In a strict vegetarian diet, antioxidant supplementation would therefore be likely covered by the normal daily intake of cisgenic tomatoes, but the levels of BCAAs would be insufficient in this single product. To be effective, a similar approach could be implemented in parallel in other fruit and vegetables, particularly if increases in free amino acids were also reflected in increases in amino acid composition in proteins in the vegetarian diet.

Multiple biofortification of crops has been earlier successfully achieved using transgenic approaches (Díaz-Gómez et al., 2017; Zhu et al., 2013). The tomato fruits described in this work are an example of how new breeding techniques can also be applied to customize food composition and adapt it to new needs that arise in modern societies, especially in the context of a steady reduction of the intake of meat-derived products for sustainability and/or ethical reasons. The increasing availability of crop genomic data will not only help us to understand crop genetics but also will provide us with new, well-defined DNA components that will enable cisgenic engineering of crop metabolism with increasing accuracy and safety.

### Experimental procedures

#### Cloning procedures

All plasmids were assembled with GoldenBraid standard procedures (Sarrion-Perdigones et al., 2013). SlALS gene (Solyc03g044330.1) was amplified from MoneyMaker genomic DNA. Three nucleotide changes were introduced on the ALS coding sequence, two of them for removing internal BsaI and BsmBI restriction sites due to cloning requirements and a third mutation that leads the Pro>Ser amino acid change at position 186 conferring CLS resistance. To introduce the mutations on the SlALS and to create level 0 parts GB0816, GB0914 and GB0144, we followed the GB domestication standard procedures (Sarrion-Perdigones et al., 2013) with primers listed on Table S4. GB0080, GB0142 (Sarrion-Perdigones et al., 2013) and GB0075 (Vazquez-Vilar et al., 2015) had been adapted to the GB standard in previous works. Level 0 parts making the selection marker transcriptional unit (GB0080, GB0816 and GB0142) were assembled in the pDGB1α1R GBvector with a restriction-ligation reaction to create the level 1 transcriptional unit GB0818. In the same way, the MYB12 fruit-specific expression-cassette was assembled in the pDGB1α2 GBvector from Level 0 parts GB0075, GB0914 and GB0144. Finally, the selection marker and the MYB12 fruit-specific expression-cassette were combined in a binary reaction in the pDGB3Ω1 vector generating level>1 element GB0830.

#### Tomato transformation

GB0830 was transferred to *Agrobacterium tumefaciens* LBA4404 strain for stable tomato transformation. Tomato (var. MoneyMaker) transformation was carried out as described by Ellul et al. (2003) with minor modifications. Briefly, cotyledons of 10 days tomato plants were cut and explants were submerged in the Agrobacterium culture for half an hour. Next, they were transferred to coculture medium and kept in the dark for 48 hours. Then, explants were transferred to the organogenesis medium with different doses of chlorsulfuron. Chlorsulfuron (purchased from Chemservice, N-11461-100 mg) was resuspended in a small volume of 1M KOH and diluted to the desired final concentration with distilled water. Individual shoots were excised and transferred to elongation medium prior to be transferred to rooting medium for root regeneration.

#### Plant material

Wild-type and T2 and T3 AM plants were grown under natural light and controlled temperature conditions (24 ºC during the day, 18 ºC at night) in a greenhouse. For initial characterization including phenylpropanoids, volatiles and primary metabolites relative quantification, red ripe fruits were collected, frozen in liquid nitrogen, ground in a cryogenic mill and stored at -80 ºC until analysis. For RT-PCR, RNA-seq, untargeted metabolomics and absolute flavonol and BCAAs quantification, fruit were checked daily after reaching the mature green stage and collected at different days post breaker depending on the analysis. For all the analyses except RT-PCR, the peel of three to five fruits was separated and pericarp and peel were independently frozen in liquid nitrogen, ground and stored at - 80 ºC until analysis.

#### Determination of ALS activity in leaves

The determination of the ALS activity was performed as described in Shimizu et al., (2008) with minor modifications. Tomato leaf discs (d=1,5 cm) of 2 months-old plants were excised and incubated on plates with 25% MS medium containing 0,5 mM 1,1-cyclopropanedicarboxylic acid and with or without 50 µg / L chlorsulfuron. After 42 hours, samples were frozen in liquid nitrogen and subsequently homogenized and extracted with 200 µl of 0.025% Triton X-100 followed by 15 min of centrifugation (12000 x*g*). Next steps were performed as indicated in Shimizu et al., (2008).

#### Determination of primary metabolites

The relative levels of polar metabolites were determined as described in (Zanor et al., 2009). One hundred mg of frozen tomato fruit powder were extracted in 1.4 mL of methanol and 60 µL of an aqueous solution with 0.2 mg mL^−1^ of ribitol, which was used as internal standard. Extraction was performed at 70 ºC for 15 min in a water bath. The extract was centrifuged at 14,000 rpm for 10 min, and the supernatant was recovered and fractionated adding chloroform and Milli-Q water. After vigorous vortexing and centrifugation at 4,000 rpm for 15 min, 50 µL of the aqueous phase were recovered and dried overnight in a speed-vac. The dry residue was subjected to a double derivatization procedure with methoxyamine hydrochloride (20 mg mL^−1^ in pyridine, Sigma) and *N*-Methyl-*N*-(trimethylsilyl)trifluoroacetamide (Macherey-Nagel). Fatty acid methyl esters (C8_-C_24) were added and used as retention index (RI) markers. Analyses were performed on a 6890N gas chromatograph (Agilent Technologies) coupled to a Pegasus 4D TOF mass spectrometer (LECO). Chromatography was performed with a BPX35 (30 m, 0.32 mm, 0.25 µm) capillary column (SGE Analytical Science) with a 2 mL min^−1^ constant helium flow. Oven programming conditions were as follows: 2 min of isothermal heating at 85 °C, followed by a 15 °C min^−1^ temperature ramp up to 360 °C. Injection temperature was set at 230 °C, and the ion source was adjusted to 250 °C. Data were acquired after electron impact ionization at 70 eV, and recorded in the 70–600 m/z range at 20 scans s^−1^. Chromatograms were analyzed by means of the ChromaTOF software. Metabolites were identified by comparison of both mass spectra and retention time with those of pure standards injected under the same conditions. Peak area of each identified compound was normalized to the internal standard area (ribitol) and sample dry weight. Five replicates per line were performed.

#### Absolute quantification of valine, leucine and isoleucine

For amino acid quantification, 100 mg of tomato fruit samples were homogenized with liquid nitrogen and extracted in 1400 µl 100% methanol supplemented with 60 µL internal standard (0.2 mg/mL ribitol). After extraction for 15 min at 70 °C and centrifugation for 10 min at 14,000 rpms, 500 µl of the supernatant were transferred to a new vial. Then 250 µl of _CHCl_3 and 500 µl of water were added. The mixture was vortexed for 15 s and centrifuged for 15 min at 14,000 rpms. 110 µl aliquots of the aqueous phase were speed-dried for 3 hours. For derivatisation, dry residues were resuspended in 40 µl of 20 mg/ml methoxyamine hydrochloride in pyridine and incubated for 90 min at 37 °C. Then, 70 µL MSTFA (N-methyl-N-[trimethylsilyl]trifluoroacetamide) and 6 µl of a retention time standard mixture (3.7% [w/v] mix of fatty acid methyl esters ranging from 8 to 24C) were added and the samples were incubated for 30 min at 37 °C.

Sample volumes of 2 µl were injected in 1:10 split mode in a 6890 N gas chromatograph (Agilent Technologies Inc. Santa Clara, CA) coupled to a Pegasus4D TOF mass spectrometer (LECO, St. Joseph, MI). Gas chromatography was performed on a BPX35 column (30 m × 0.32 mm × 0.25 μm) (SGE Analytical Science Pty Ltd., Australia) with helium as carrier gas, constant flow 2 mL min^−1^. The liner was set at 250 °C. Oven program was 85 °C for 2 min, 8 °C min^−1^ ramp until 360 °C. Mass spectra were collected at 6.25 spectra s^―1^ in the m/z range 35–900 and ionization energy of 70 eV. Chromatograms and mass spectra were evaluated using the CHROMATOF program (LECO, St. Joseph, MI). For metabolite identification and absolute quantification a standard curve was made with authentic standards.

#### Determination of volatile compounds

Volatile compounds were captured by means of headspace solid phase microextraction (HS-SPME) and separated and detected by means of gas chromatography coupled to mass spectrometry (GC/MS). Samples were processed as described in Rambla et al. (2017) with minor modifications. Roughly, 500 mg of frozen tomato powder were introduced in a 15 mL glass vial, the cap closed and incubated at 37 °C for 10 min in a water bath. 500 mL of an EDTA 100 mM, pH 7.5 solution and 1.1 g of _CaCl_2_.2H_2O were added, gently mixed and sonicated for 5 min. One mL of the resulting paste was transferred to a 10 mL screw cap headspace vial with silicon/PTFE septum and analyzed within 10 hours. Volatile compounds were extracted from the headspace by first preincubating the vials at 50 °C for 10 min under 500 rpm agitation. A 65 µm PDMS/DVB SPME fiber (SUPELCO) was then introduced in the vial and exposed to the headspace for 20 min, with identical conditions of agitation and temperature. The volatile compounds adsorbed in the fiber were desorbed in the injection port of the gas chromatograph at 250 °C for 1 min in splitless mode. The fiber was then cleaned at 250 °C for an additional 5 min in an SPME Fiber Conditioning Module (CTC Analytics) to prevent cross-contamination between samples. Incubation, extraction, injection and fiber cleaning were performed by means of a CombiPAL autosampler (CTC Analytics). Chromatography was performed on a 6890N gas chromatograph (Agilent) with a DB-5ms (60 m, 0.25 mm, 1.00 µm) capillary column (J&W) with a constant helium flow of 1.2 mL min^−1^. Oven ramp conditions were: 40 °C for 2 min, 5 °C min^−1^ ramp until 250 °C and a final hold at 250 °C for 5 min. GC interface and MS source temperatures were 260 °C and 230 °C respectively. Detection was performed in a 5975B mass spectrometer (Agilent) in the 35–300 m/z range at 6.2 scans/s, with 70 eV electron impact ionization. Data were recorded and processed with the Enhanced ChemStation E.02.02 software. Unequivocal identification of volatile compounds was performed by comparison of both mass spectra and retention time with those of pure standards (Sigma). For quantitation, one specific ion was selected for each compound and the corresponding peak from the extracted ion chromatogram was integrated. An admixture reference sample was prepared for each season by mixing thoroughly equal amounts of each sample. A 500 mg aliquot of the admixture was analyzed every six samples and processed as any other sample as part of the injection series. This admixture was used as a reference to normalize for temporal variation and fiber aging.

#### LC-ESI(+/-)-MS analysis of tomato fruit phenylpropanoids

Phenylpropanoid extraction was carried out as previously described (Fasano et al., 2016). Briefly, 10 mg of ground freeze-dried fruit powder were extracted with 0.75 mL cold 75% (v/v) methanol, 0.1% (v/v) formic acid, spiked with 10 µg mL^−1^ formononetin. Samples were vortexed for 30 s, and shaken for 15 min at 15 Hz using a Mixer Mill 300 (Qiagen) and kept at RT for 5 min (twice). After centrifugation for 15 min at 20,000 x*g* at 4 °C, 0.6 mL of supernatant were removed and transferred to HPLC tubes. For each genotype, extractions from 3 fruits were performed. Separation was carried out using an Ultimate 3000 HPLC coupled to a Q-EXACTIVE mass spectrometer (ThermoFisher) equipped with a C18 Luna reverse-phase column (150 × 2.0 mm, 3 µm; Phenomenex, Macclesfield, UK) and a gradient system as follows: 95%A:5%B for one minute, followed by a linear gradient to 25%A:75%B over 40 minutes. LC conditions were kept for 2 more minutes, before going back to the initial LC conditions in 18 min. 10 µl of each sample were injected and a flow of 0.2 mL was carried out during the whole LC runs. Detection was performed continuously from 230 to 800 nm with an online Ultimate 3000 photodiode array detector (PDA, Thermo Fischer Scientific, Waltham, MA). All solvents used were LC-MS grade quality (CHROMASOLV^®^ from Sigma-Aldrich). The Exactive Plus Orbitrap mass spectrometer was equipped with a heated electrospray probe (H-ESI). ESI and MS parameters were as follows: spray voltage ―5.0 kV, sheath gas and auxiliary nitrogen pressures 30 and 10 arbitrary units, respectively; capillary and heater temperatures were set at, respectively 250 and 150 °C, while tube lens voltage was 50 V. Data were acquired in profile mode. Identification was performed as reported before (Diretto et al., 2020), through comparison of chromatographic and spectral properties of authentic standards and reference spectra, and on the basis of the m/z accurate masses, as reported on Pubchem database (http://pubchem.ncbi.nlm.nih.gov/) for monoisotopic masses identification, or on Metabolomics Fiehn Lab Mass Spectrometry Adduct Calculator (http://fiehnlab.ucdavis.edu/staff/kind/Metabolomics/MS-Adduct-Calculator/) in case of adduct ion detection. Metabolites were relatively quantified on the basis of the internal standard amounts. ANOVA was applied to the absolute metabolite quantifications, and pair-wise comparisons were used in Tukey’s HSD test running on the Past 3.x software (http://folk.uio.no/ohammer/past/).

#### Flavonol absolute quantification

50 mg of frozen tomato grinded tissue were resuspended in 500 µl of 75% acetonitrile with 1 ppm genistein. The homogenate was vortexed, sonicated for 10 min and centrifuged for 5 min at 14,000 rpms. 200 µl of the supernatant were mixed with an equal volume of HCl 2N and incubated at 80 °C for 2 hours. Samples were filtered with a 0.2 µM filter and 1 µl was injected. Three replicates of each genotype were extracted and analysed. Flavonol quantification was performed using a Orbitrap Exploris 120 mass spectrometer coupled with a Vanquish UHPLC System (Thermo Fisher Scientific, Waltham, MA, USA). LC was carried out by reverse-phase ultraperformance liquid chromatography using a Acquity PREMIER BEH C18 UPLC column (1.7 µM particle size, dimensions 2.1 × 150 mm) (Waters Corp., Mildford, MA, USA). The mobile phase consisted 0.1% formic acid in water (phase A), and 0.1% formic acid in acetonitrile (phase B). The solvent gradient program was conditioned as follows: 0.5% solvent B over the first 2 min, 0.5–30% solvent B over 25 min, 30–100% solvent B over 13 min, 2 min at 100% B, return to the initial 0.5% solvent B over 1 min, and conditioning at 0.5% B for 2 minutes. The flow rate was 0.4 mL min^−1^ and the injection volume was 1 µl. The column temperature was set at 40 °C. Ionisation was performed with heated electrospray ionization (H-ESI) in positive and negative mode. Samples were acquired in full scan mode (resolution set at 120000 measured at FWHM). For absolute quantification a standard curve was performed with authentic standards using genistein as internal standard. Data processing was performed with TraceFinder software (Thermo Scientific, Waltham, MA, USA).

#### LC-MS untargeted analysis of polar extracts

Flesh and peel samples of five fruits collected at 4 and 7 dpb were frozen in liquid nitrogen and ground into a fine powder. Next, samples were freeze-dried and weighed (10 ± 0.5 mg) and extracted in methanol/water (1:1) as described previously. To each aliquot of the polar phase (100 µl), internal standard (genistein, 1 µg) was added. The polar phase was used for metabolite profiling by UPLC-ESI-QToF 6560 (Agilent Technologies, Stockport, UK) as previously described (Drapal et al., 2022). Data processing was performed with Agilent Profinder (v10.0 SP1; Agilent Technologies, Inc.). The identified molecular features were relatively quantified to the internal standard and dry weight of the sample. Only molecular features that were present in at least two out of the five biological replicates for either AM or WT were analyzed. PCA and hierarchical clustering analysis were performed with Metaboanalyst using auto scaling for data normalization.

#### RNA isolation, RT-qPCR and RNA sequencing

For RT-qPCR analyses, fruits from AM plants were harvested at different ripening stages and frozen and ground using liquid nitrogen. Total RNA was extracted with TRIzolTM Reagent following manufacturer instructions. 4 µg of total RNA was treated with a DNase-I Invitrogen Kit following manufacturer’s instructions. Then, 1 µg of treated RNA was used for cDNA synthesis using a PrimeScript™ RT-PCR Kit (Takara, Kusatsu, Japan) in a final volume of 20 μl. Als and Myb12 expression levels were measured in three technical replicates reactions in the presence of a fluorescent dye (SYBR® Premix Ex Taq) using QuantStudio 3 equipment (Applied Biosystems, Waltham, MA, USA). The specific primers for detection of endogenous and cisgenic AlS and Myb12 genes are listed in Table S1. Actin was used as an internal reference gene. Calculations were carried out according to the comparative ΔΔCt method.

For transcriptomic analysis, both WT and AM fruits were marked at breaker and fruits were collected 4 days after breaker. Fruit flesh was frozen and ground using liquid nitrogen. Total RNA was extracted with TRIzolTM Reagent following manufacturer instructions. Both library preparation and RNA-seq sequencing were performed by Novogene (China) with total RNA, generating Paired end (PE) 2×150bp reads with NovaSeq 6000 PE150 platform.

#### RNA-seq preprocessing and expression profiles analysis

Sequence reads were quality checked using FastQC v0.11.9 (https://www.bioinformatics.babraham.ac.uk/projects/fastqc/). Raw reads were quality trimmed and Illumina adaptors were removed with Trimmomatic v0.39 (Bolger et al., 2014). Clean reads were mapped against *Solanum lycopersicum* reference genome SL4.0 (Hosmani et al., 2019) using Hisat2 v2-2.2.1 (Kim et al., 2019) and gene abundances were calculated using StringTie v2.1.6 (Pertea et al., 2015) with the genome annotation ITAG4.0 (Hosmani et al., 2019). From these counts a gene expression table of raw read counts was generated. Raw gene expression was normalized by trimmed mean of M-values and differential expression analysis was performed using edgeR v3.36.0 with a negative binomial exact test using both common and tagwise dispersion (Robinson et al., 2010). Then, significant differential expressed genes were kept if their false discovery rate (FDR) was less than 0.05 and their average expression for all samples was equal or greater than 1 count per million mapped (CPM). Kept genes were mapped against KEGG terms database using blastkoala (Kanehisa et al., 2016). The number of reads corresponding to endogenous and cisgenic MYB12 and ALS was calculated using single nucleotide differences by allele counting using samtools mpileup v1.7 (H. Li, 2011).

## Data availability

RNA-Seq data reported in this paper is available at NCBI database (Bioproject PRJNA821387).

## Funding

This work has been funded by grant PID2019-108203RB-100 from the Spanish Ministerio de Ciencia e Innovación, through the Agencia Estatal de Investigación (co-financed European Regional Development Fund). MVV is recipient of APOSTD/2020/096 (Generalitat Valenciana and Fondo Social Europeo post-doctoral grant). JLR acknowledges support by the Spanish Ministry of Science and Innovation through a *Juan de la Cierva-Incorporación* grant (IJC2020-045612-I).

## Conflict of interest

The authors declare that they have no conflict of interest.

### List of Supplementary Material

**Figure S1**. Colorimetric assay for the determination of ALS activity in leaf extracts of three independent T2 AM lines and their corresponding azygous lines in the absence and presence of 50 µg/L chlorsulfuron. Red indicates ALS activity.

**Figure S2**. Overexpression of the acetolactate synthase gene results in increased branched chain amino acids in leaves.

**Figure S3**. Overexpression of the acetolactate synthase gene results in unintended increased branched chain volatiles in tomato fruits.

**Figure S4**. Expression levels of endogenous and cisgenic ALS and MYB12 genes in tomato fruit in different tissues.

**Figure S5**. Fruit specific overexpression of the SlMyb12 gene results in increased phenylpropanoids in tomato fruits.

**Figure S6**. Transcriptomic analysis of ALS-MYB flesh samples shows overexpression of multiple genes involved in the amino acids biosynthetic pathway.

**Table S1**. List of differentially expressed genes between AM and WT flesh samples.

**Table S2**. List of GO terms with top significance retrieved from the RNAseq analysis of the AM fruits.

**Table S3**. Top 25 genes overexpressed in the AM fruits contributing to the chloroplast and plastid-related GO terms of the cellular component.

**Table S4**. List of primers used in this work.

## Notes

### Competing Interest Statement

The authors have declared no competing interest.

