## Supplementary Information for "Dually biofortified cisgenic tomatoes with increased flavonoids and branched-chain amino acids content"

### Supporting Information

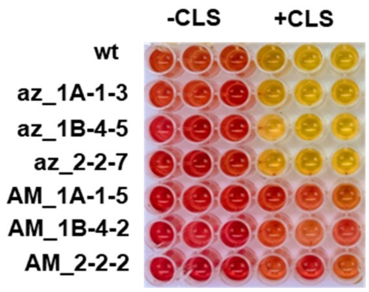

**Figure S1.** Colorimetric assay for the determination of ALS activity in leaf extracts of three independent T2 AM lines and their corresponding azygous lines in the absence and presence of 50 µg/L chlorsulfuron. Red indicates ALS activity.

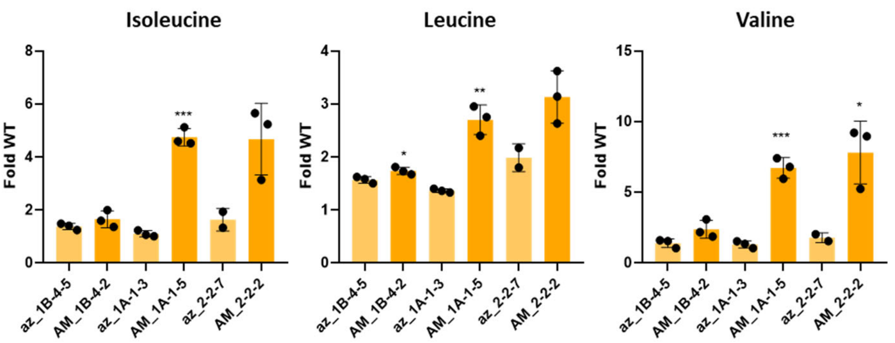

**Figure S2.** Overexpression of the acetolactate synthase gene results in increased branched chain amino acids in tomato leaves.

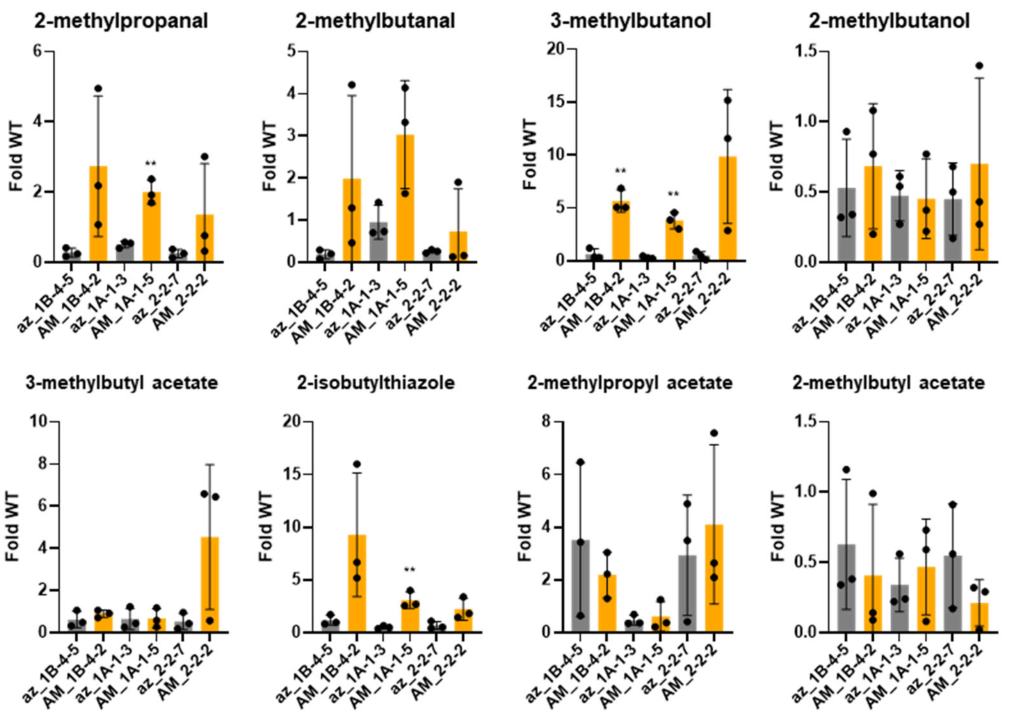

**Figure S3.** Overexpression of the acetolactate synthase gene results in unintended increase of most branched chain volatiles in tomato fruits.

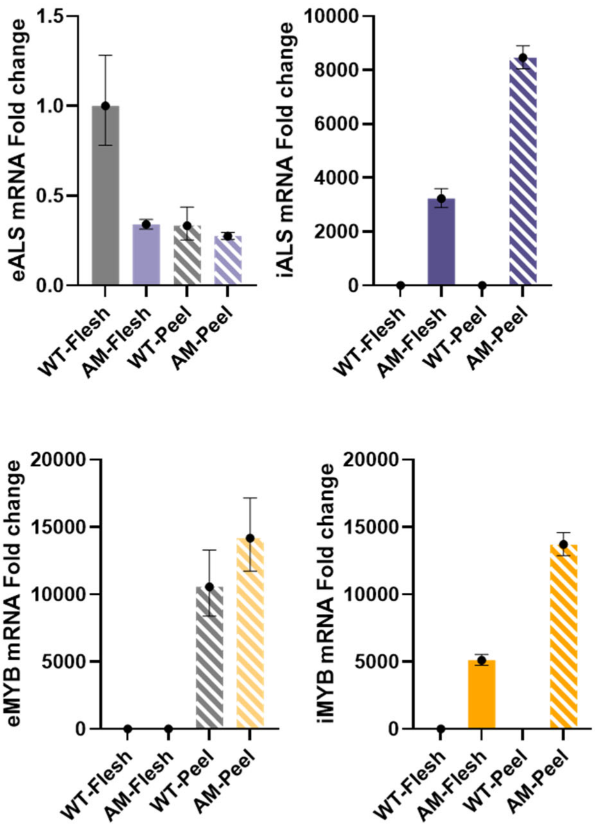

**Figure S4.** Expression levels of endogenous and intragenic ALS and MYB12 genes in tomato fruit in different tissues.

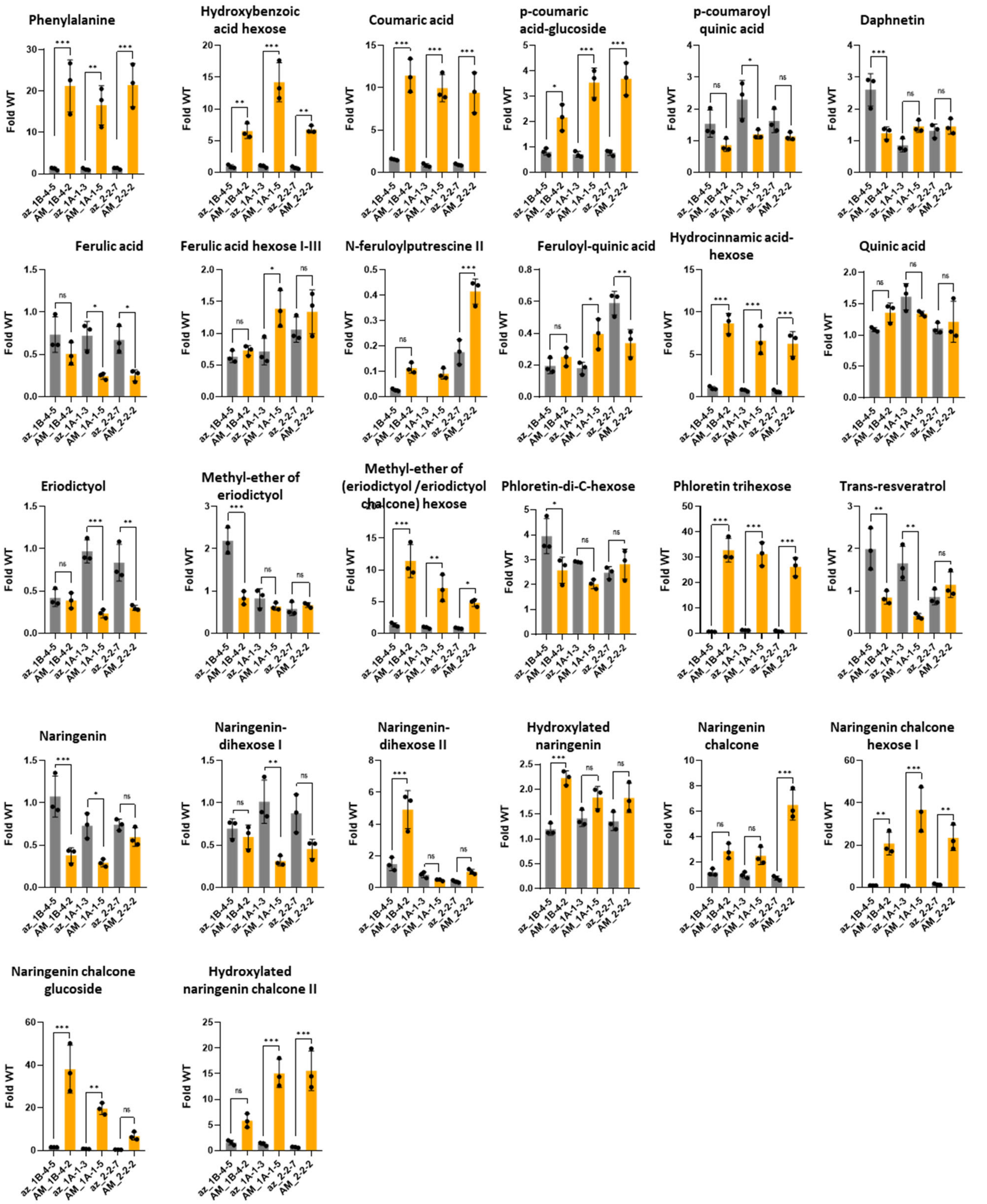

**Figure S5.** Fruit specific overexpression of the SlMyb12 gene results in increase of most phenylpropanoids in tomato fruits.

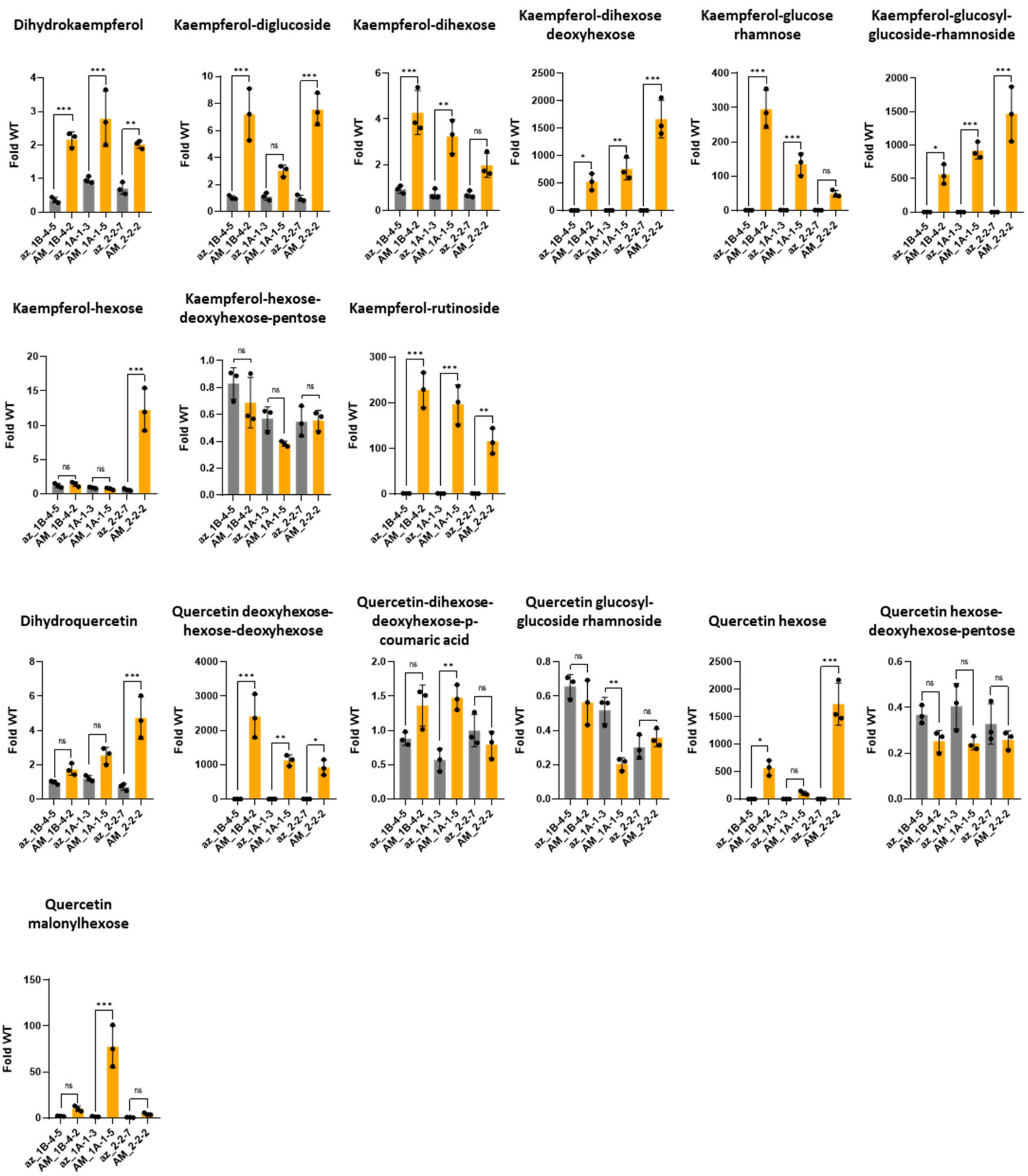

**Figure S5 (continuation).**

A

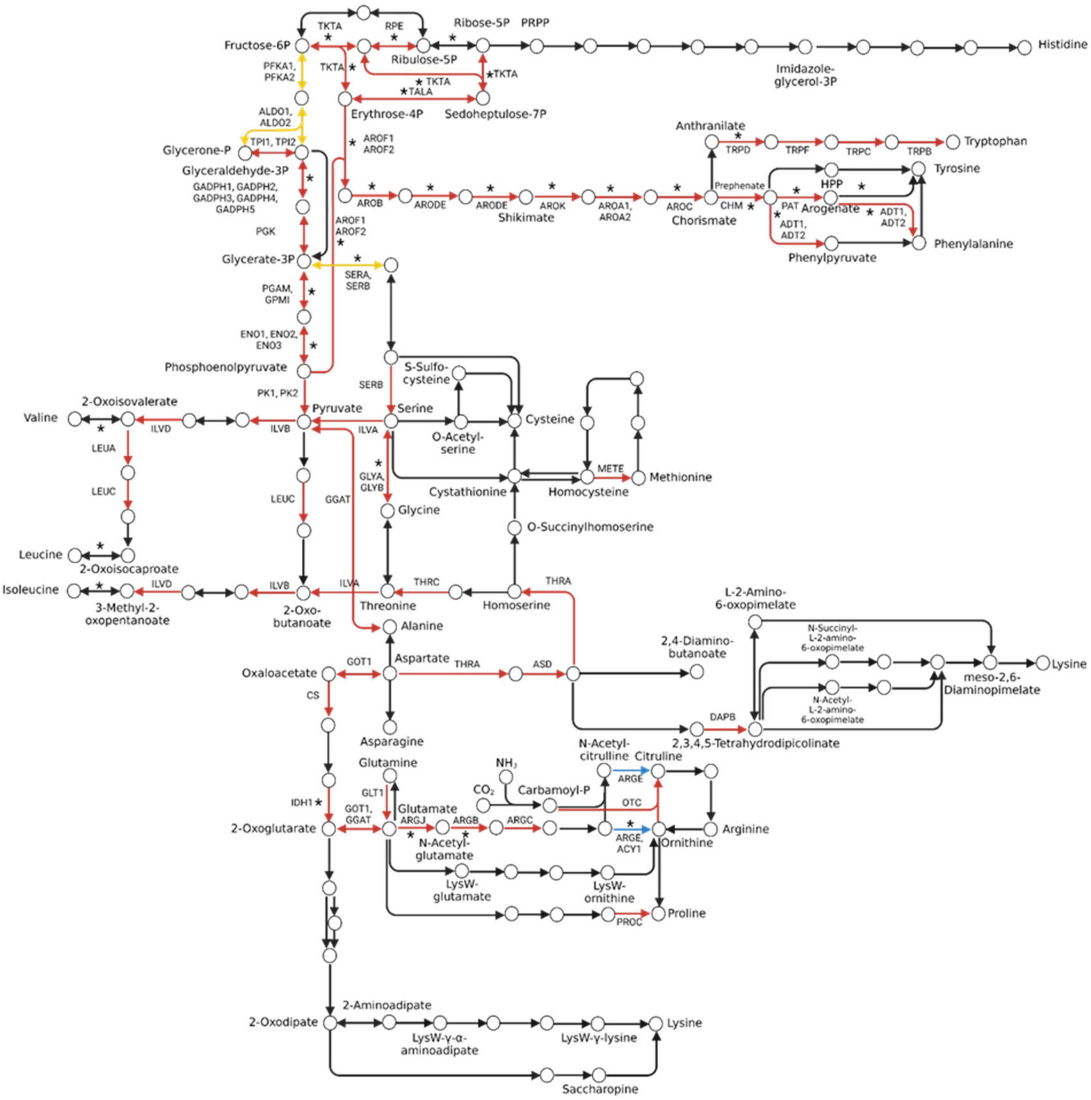

**Figure S6. Transcriptomic analysis of ALS-MYB flesh samples shows overexpression of multiple genes involved in the amino acids biosynthetic pathway.** A) A KEGG analysis shows that several genes of the amino acids biosynthetic pathway (red arrows) are overexpressed in the AM fruits compared to WT. Asterisks indicate genes also overexpressed in AtMYB12 fruits from Zhang et al. (2015). B) Log FPKMs of the genes differentially expressed among lines in the amino acids biosynthesis pathway depicted in A).

B

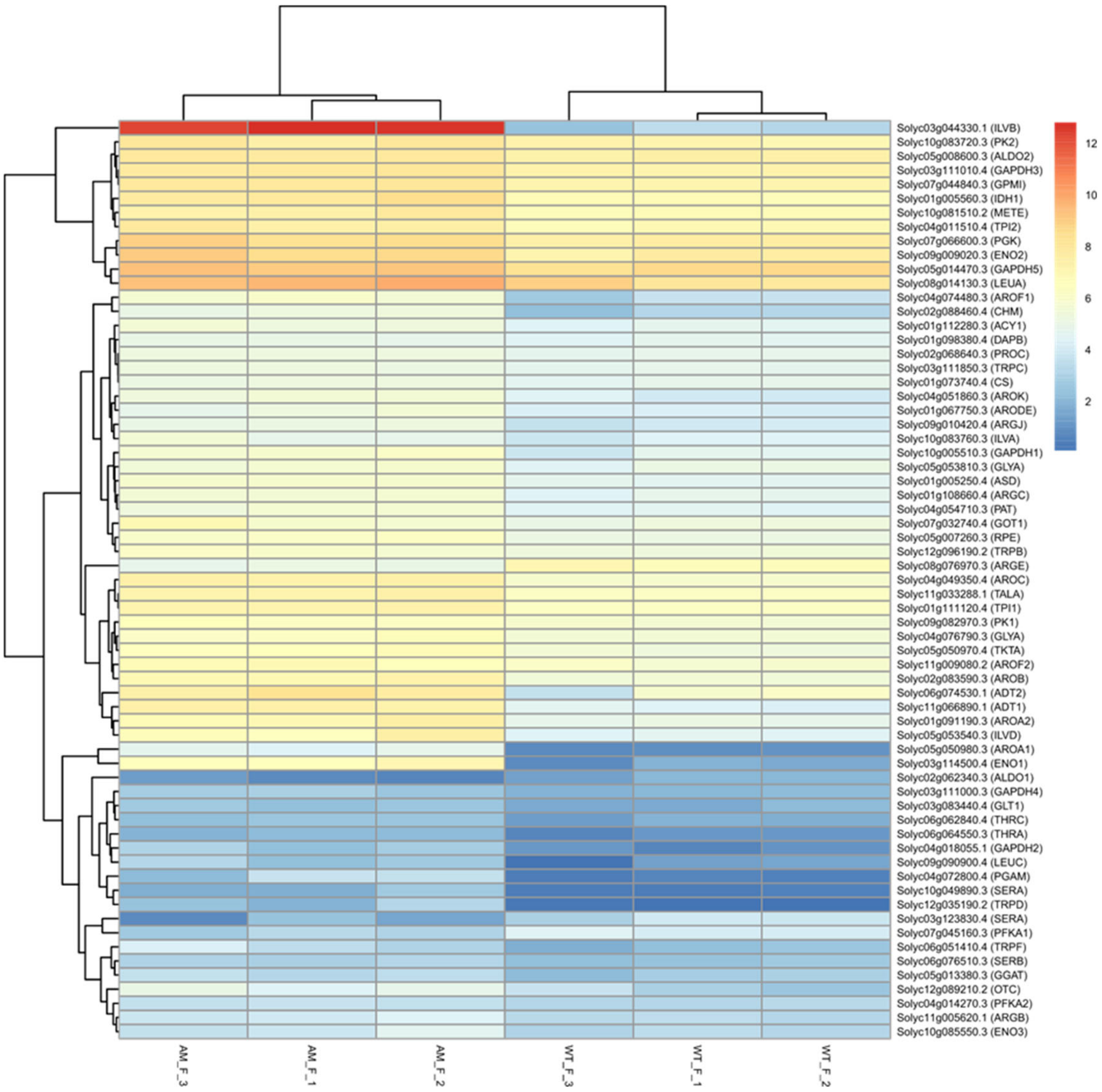

**Figure S6 (continuation).**

**Table S1. List of differentially expressed genes between AM and WT flesh samples.**

| gene | description | KEGG | logFC | FDR | AM_F_1 | AM_F_2 | AM_F_3 | WT_F_1 | WT_F_2 | WT_F_3 |
|---|---|---|---|---|---|---|---|---|---|---|
| Solyc09g059020.4 | Quinone-oxidoreductase QR1, chloroplastic (AHRD V3.3 *** QR1_TRIVS) | CEQORH | -9.275 | 7.00E-197 | 55.625 | 45.481 | 55.308 | 0.135 | 0.063 | 0.039 |
| Solyc03g044330.1 | Acetolactate synthase (AHRD V3.3 *** C0L093_TOBAC) | ILVB | -9.673 | 1.89E-92 | 7182.733 | 6395.479 | 4924.452 | 10.347 | 8.270 | 4.438 |
| Solyc09g059060.1 | Quinone-oxidoreductase QR1, chloroplastic (AHRD V3.3 *-* QR1_TRIVS),Pfam:PF13602 | NA | -12.768 | 4.29E-73 | 72.393 | 64.770 | 45.432 | 0.000 | 0.000 | 0.000 |
| Solyc05g050980.3 | 3-phosphoshikimate 1-carboxyvinyltransferase (AHRD V3.3 *** A0A0M8KSM3_NICAT) | AROA | -4.822 | 2.05E-70 | 19.820 | 29.011 | 26.514 | 0.977 | 1.012 | 0.702 |
| Solyc11g066890.1 | Arogenate dehydratase (AHRD V3.3 *** A0A2G3BBC7_CAPCH) | ADT | -3.002 | 4.00E-49 | 168.999 | 153.605 | 162.419 | 19.368 | 18.481 | 22.696 |
| Solyc09g059070.3 | Quinone-oxidoreductase chloroplastic-like (AHRD V3.3 *-* A0A2K3P901_TRIPR) | NA | -11.991 | 3.95E-48 | 27.090 | 21.026 | 14.324 | 0.000 | 0.000 | 0.000 |
| Solyc06g075010.4 | 60 kDa chaperonin (AHRD V3.3 *** B2IXD2_NOSP7) | NA | -5.960 | 9.48E-47 | 33.751 | 45.950 | 38.851 | 0.837 | 0.789 | 0.314 |
| Solyc10g084400.2 | Glutathione S-transferase (AHRD V3.3 *** Q76KW1_PEA) | GST | -2.941 | 9.92E-43 | 171.003 | 144.622 | 129.719 | 16.676 | 20.701 | 20.768 |
| Solyc08g067310.1 | Non-specific serine/threonine protein kinase (AHRD V3.3 *** A0A2G2W7U9_CAPBA) | NA | -4.071 | 2.74E-38 | 7.858 | 11.653 | 11.531 | 0.575 | 0.751 | 0.525 |
| Solyc04g011390.1 | Histone H4 (AHRD V3.3 *-* F2E7L1_HORVV) | H4 | -4.686 | 2.17E-37 | 29.441 | 22.535 | 33.298 | 1.413 | 1.223 | 0.641 |
| Solyc01g008920.4 | NAD(P)-binding Rossmann-fold superfamily protein (AHRD V3.3 *** A0A2U1L7S8_ARTAN) | NA | -2.977 | 1.35E-30 | 15.560 | 13.381 | 17.695 | 1.847 | 1.732 | 2.321 |
| Solyc06g074790.2 | Histone H2B (AHRD V3.3 *** A0A2G3CAQ8_CAPCH) | H2B | -3.575 | 5.87E-26 | 33.634 | 30.532 | 47.391 | 3.180 | 4.178 | 2.105 |
| Solyc10g007100.3 | Protein DETOXIFICATION (AHRD V3.3 *** A0A218WWW8_PUNGR) | TC.MATE | -3.341 | 1.20E-25 | 15.783 | 13.645 | 21.278 | 2.291 | 1.472 | 1.277 |
| Solyc03g114500.4 | Enolase (AHRD V3.3 *** A0A2G2WR88_CAPBA) | ENO | -5.816 | 2.97E-25 | 91.658 | 135.238 | 93.086 | 2.706 | 2.379 | 0.746 |

| | | | | | | | | | | |
|---|---|---|---|---|---|---|---|---|---|---|
| Solyc03g025840.3 | Cytochrome b561/ferric reductase transmembrane protein family (AHRD V3.3 *** A0A178UZ09_ARATH) | CYB561 | -2.894 | 1.15E-23 | 386.716 | 532.787 | 343.148 | 51.572 | 53.732 | 65.298 |
| Solyc06g083440.3 | Cytochrome b5 (AHRD V3.3 *** A0A2G3CCB6_CAPCH) | CYB5 | -2.895 | 2.07E-22 | 8.456 | 9.204 | 9.286 | 0.976 | 1.584 | 1.062 |
| Solyc06g073190.3 | fructokinase 2 | SCRK | -1.687 | 9.22E-22 | 141.859 | 115.105 | 141.035 | 42.170 | 44.897 | 37.392 |
| Solyc10g083360.2 | Calmodulin-binding family protein, putative, expressed (AHRD V3.3 *** Q2QXN6_ORYSJ) | NA | -1.778 | 1.06E-20 | 87.607 | 82.757 | 73.546 | 26.617 | 26.046 | 19.271 |
| Solyc06g005160.4 | cytosolic ascorbate peroxidase 1 | L-ascorbate peroxidase | -1.731 | 5.71E-19 | 580.554 | 662.528 | 647.435 | 191.002 | 191.157 | 191.100 |
| Solyc06g005430.1 | Histone H4 (AHRD V3.3 *-* F2E7L1_HORVV) | H4 | -1.903 | 8.15E-19 | 45.626 | 40.983 | 45.468 | 11.450 | 13.942 | 10.237 |
| Solyc10g005300.3 | Serine/threonine-protein kinase PBS1 (AHRD V3.3 *** A0A2G2VSF8_CAPBA) | NA | -3.347 | 1.55E-18 | 1.860 | 2.165 | 1.846 | 0.151 | 0.262 | 0.151 |
| Solyc05g053100.3 | Dihydrolipoyl dehydrogenase-like protein (AHRD V3.3 *** A0A2K3PDU3_TRIPR) | DLD | -2.126 | 6.86E-18 | 5.278 | 4.385 | 3.899 | 1.084 | 1.027 | 1.005 |
| Solyc09g065180.3 | binding protein precursor AF106660 | NA | -2.061 | 3.65E-17 | 7.585 | 8.886 | 8.732 | 1.940 | 1.879 | 2.238 |
| Solyc11g056680.1 | Leucine-rich repeat receptor-like protein (AHRD V3.3 *** H6V788_MALDO) | NA | -2.784 | 6.79E-17 | 27.971 | 15.487 | 22.810 | 2.716 | 2.867 | 3.957 |
| Solyc04g049350.4 | chorismate synthase 1 precursor | AROC | -1.543 | 1.34E-16 | 157.900 | 174.989 | 166.767 | 56.848 | 57.173 | 58.446 |
| Solyc04g055170.3 | annexin p35 | ANNAT | -1.557 | 2.88E-16 | 63.446 | 58.220 | 59.447 | 22.294 | 24.195 | 15.867 |
| Solyc01g091190.3 | 5-enolpyruvylshikimate-3-phosphate synthase | AROA | -2.242 | 3.92E-16 | 123.963 | 183.561 | 113.263 | 32.912 | 28.620 | 28.353 |
| Solyc04g081400.3 | plastidic hexokinase | HK | -1.640 | 8.47E-16 | 15.867 | 19.020 | 16.842 | 6.508 | 5.606 | 4.675 |
| Solyc04g074480.3 | DAHP synthase 2 precursor | AROF | -2.481 | 1.09E-15 | 62.575 | 50.191 | 51.149 | 11.821 | 12.315 | 5.747 |
| Solyc01g006280.3 | Formate--tetrahydrofolate ligase (AHRD V3.3 *** A0A2G3DED7_CAPCH) | FHS | -1.853 | 3.70E-15 | 151.164 | 170.898 | 149.560 | 35.669 | 46.968 | 48.434 |
| Solyc01g099340.3 | Protein indeterminate-domain 12 (AHRD V3.3 *** A0A2G2W7U4_CAPBA) | NA | -2.176 | 6.39E-15 | 26.801 | 24.458 | 38.969 | 7.264 | 8.109 | 4.859 |
| Solyc02g088460.4 | Chorismate mutase (AHRD V3.3 *** B5LAU1_CAPAN) | Chorismate mutase | -2.464 | 9.53E-15 | 38.417 | 41.051 | 32.298 | 8.079 | 8.612 | 3.969 |
| Solyc06g005150.3 | ascorbate peroxidase | L-ascorbate peroxidase | -1.436 | 1.37E-14 | 227.118 | 230.826 | 266.343 | 96.013 | 88.414 | 85.081 |

| | | | | | | | | | | |
|---|---|---|---|---|---|---|---|---|---|---|
| Solyc06g005420.1 | Histone H4 (AHRD V3.3 *-* F2E7L1_HORVV) | H4 | -1.676 | 1.92E-14 | 124.996 | 93.953 | 117.118 | 34.692 | 31.472 | 38.851 |
| Solyc04g082460.4 | Catalase (AHRD V3.3 *** A0A2G2X1I6_CAPBA) | KATE | -1.598 | 4.18E-13 | 57.532 | 42.834 | 47.139 | 13.867 | 16.592 | 18.245 |
| Solyc06g065210.4 | Neutral/alkaline invertase (AHRD V3.3 *** I0CL58_MANES) | NA | -1.458 | 5.49E-13 | 10.476 | 10.727 | 12.320 | 3.816 | 4.861 | 3.634 |
| Solyc05g053540.3 | Dihydroxy-acid dehydratase (AHRD V3.3 *** A0A2G3CHU1_CAPCH) | ILVD | -2.524 | 2.24E-12 | 85.893 | 177.627 | 96.939 | 22.574 | 21.844 | 19.227 |
| Solyc09g018510.3 | Phytol kinase (AHRD V3.3 *** A0A2K3MZ75_TRIPR) | FOLK | -1.346 | 3.09E-12 | 17.558 | 15.176 | 17.548 | 7.346 | 7.099 | 5.496 |
| Solyc02g088340.4 | WRKY transcription factor 3 | NA | -3.005 | 4.42E-12 | 155.654 | 113.589 | 198.823 | 24.012 | 25.775 | 9.687 |
| Solyc06g005260.3 | Glutaredoxin (AHRD V3.3 *** A0A2G2XHT5_CAPBA) | GRXC | -1.435 | 4.87E-12 | 290.086 | 339.641 | 273.136 | 120.519 | 127.524 | 90.546 |
| Solyc02g083590.3 | dehydroquinate synthase | AROB | -1.511 | 8.45E-12 | 117.171 | 151.383 | 114.395 | 47.951 | 43.502 | 44.055 |
| Solyc09g072590.3 | actin-depolymerizing factor (AHRD V3.3 *** A0A2I4EMY6_9ROSI) | CFL | -1.303 | 1.62E-11 | 101.897 | 129.610 | 117.086 | 47.440 | 53.823 | 41.706 |
| Solyc10g086580.2 | Ribulose bisphosphate carboxylase/oxygenase activase, chloroplastic (AHRD V3.3 *** RCA_SOLPN) | NA | -1.196 | 2.40E-11 | 88.758 | 82.519 | 98.850 | 40.320 | 38.802 | 39.346 |
| Solyc12g035190.2 | Anthranilate phosphoribosyltransferase (AHRD V3.3 *** A0A2U1N166_ARTAN) | TRPD | -5.191 | 3.47E-11 | 2.986 | 8.214 | 4.344 | 0.079 | 0.076 | 0.256 |
| Solyc08g066360.3 | Malic enzyme (AHRD V3.3 *** A0A2G2W5J3_CAPBA) | MAEB | -4.864 | 3.86E-11 | 7.154 | 12.025 | 2.968 | 0.206 | 0.163 | 0.381 |
| Solyc09g090700.1 | Succinate semialdehyde dehydrogenase (AHRD V3.3 *** A0A2U1MA81_ARTAN) | SSADH | -1.221 | 1.37E-10 | 89.751 | 101.619 | 102.301 | 40.615 | 41.749 | 44.209 |
| Solyc01g100930.3 | GDSL esterase/lipase (AHRD V3.3 *** A0A1U8GQF3_CAPAN) | NA | -2.799 | 1.67E-10 | 2.585 | 4.449 | 3.691 | 0.648 | 0.613 | 0.291 |
| Solyc05g006740.4 | Glutathione S-transferase (AHRD V3.3 *** A0A200QY01_9MAGN) | GST | -2.626 | 3.04E-10 | 4.029 | 6.718 | 7.570 | 1.263 | 0.780 | 0.927 |
| Solyc12g056830.1 | ATP synthase subunit delta, chloroplastic-like (AHRD V3.3 *** A0A2I4F0A5_9ROSI) | ATPF1D | -1.727 | 3.57E-10 | 32.821 | 35.387 | 54.109 | 11.115 | 13.317 | 12.698 |
| Solyc04g054710.3 | Aminotransferase (AHRD V3.3 *** A0A200QW13_9MAGN) | PAT | -1.162 | 4.06E-10 | 52.867 | 53.180 | 42.278 | 22.924 | 22.380 | 21.426 |

| | | | | | | | | | | |
|---|---|---|---|---|---|---|---|---|---|---|
| Solyc09g097960.3 | NAD(P)-linked oxidoreductase, aldo/keto reductase family protein (AHRD V3.3 *** A0A1Y1HY98_KLENI) | NA | -4.427 | 5.72E-10 | 70.634 | 45.138 | 15.137 | 1.888 | 3.167 | 1.127 |
| Solyc05g050120.3 | cytosolic NADP-malic enzyme | MAEB | -1.242 | 5.89E-10 | 195.760 | 208.265 | 187.107 | 79.485 | 83.255 | 88.299 |
| Solyc01g096430.4 | NADPH:quinone oxidoreductase-like (AHRD V3.3 *** A0A2I4GZV4_9ROSI) | CHRR | -1.589 | 7.38E-10 | 15.483 | 20.072 | 15.199 | 6.516 | 6.173 | 4.413 |
| Solyc09g007850.3 | RNA-binding protein (AHRD V3.3 *** A0A2U1PI86_ARTAN) | NA | -1.202 | 7.38E-10 | 152.336 | 115.327 | 151.359 | 60.727 | 67.051 | 55.490 |
| Solyc10g049890.3 | D-3-phosphoglycerate dehydrogenase (AHRD V3.3 *** A0A2G2VSK7_CAPBA) | SERA | -3.059 | 1.16E-09 | 2.383 | 5.610 | 2.482 | 0.382 | 0.529 | 0.361 |
| Solyc05g005760.4 | NHL domain-containing protein (AHRD V3.3 *** A0A2K3PAQ7_TRIPR) | NA | -2.175 | 1.71E-09 | 4.271 | 4.733 | 5.610 | 1.383 | 1.278 | 0.626 |
| Solyc09g010420.4 | Arginine biosynthesis bifunctional protein ArgJ, chloroplastic (AHRD V3.3 *** A0A2G2YQV3_CAPAN) | ARGJ | -1.257 | 2.15E-09 | 33.084 | 37.793 | 29.791 | 14.570 | 16.744 | 11.404 |
| Solyc09g008550.4 | NCS1 family nucleobase:cation symporter-1 (AHRD V3.3 *** A0A328X9Z8_9BURK) | TC.NCS1 | -1.912 | 3.38E-09 | 9.444 | 10.081 | 7.374 | 1.964 | 2.031 | 3.117 |
| Solyc01g088610.4 | 10 kDa chaperonin (AHRD V3.3 *** A0A2K3N899_TRIPR) | GROES | -1.081 | 3.64E-09 | 221.948 | 241.781 | 251.988 | 108.328 | 117.614 | 114.448 |
| Solyc05g007260.3 | Ribulose-phosphate 3-epimerase (AHRD V3.3 *** A0A328DMM8_9ASTE) | RPE | -1.052 | 3.73E-09 | 71.784 | 78.895 | 65.761 | 35.344 | 35.401 | 34.386 |
| Solyc04g051860.3 | shikimate kinase precursor | AROK | -1.406 | 8.63E-09 | 49.028 | 47.702 | 39.762 | 15.477 | 15.368 | 20.606 |
| Solyc09g011550.3 | Glutathione S-transferase (AHRD V3.3 *** C0LF68_CAPAN) | GST | -2.506 | 1.72E-08 | 5.384 | 2.366 | 3.965 | 0.768 | 0.720 | 0.551 |
| Solyc01g099200.3 | Lipoxygenase (AHRD V3.3 *** K4B0V7_SOLLC) | LOX1_5 | -2.808 | 2.10E-08 | 11.754 | 6.303 | 16.345 | 1.013 | 1.665 | 2.183 |
| Solyc10g005510.3 | Glyceraldehyde-3-phosphate dehydrogenase (AHRD V3.3 *** A0A2G3BCA8_CAPCH) | GAPDH | -1.526 | 3.73E-08 | 57.499 | 67.762 | 53.770 | 25.323 | 24.230 | 13.748 |
| Solyc06g060250.3 | Aldehyde dehydrogenase (AHRD V3.3 *** A0A2G2WJP7_CAPBA) | ALDH | -0.979 | 6.71E-08 | 20.109 | 16.842 | 20.305 | 10.205 | 10.664 | 8.411 |
| Solyc09g063130.3 | Photosystem I reaction center subunit IV (AHRD V3.3 *-* A0A2P6U4S6_CHLSO) | PSAE | -1.408 | 9.15E-08 | 21.128 | 24.223 | 23.499 | 7.895 | 7.383 | 10.627 |
| Solyc12g056660.3 | Mitochondrial carrier protein (AHRD V3.3 *** A0A200QJI9_9MAGN) | SLC25A26 | -1.743 | 9.21E-08 | 3.918 | 4.778 | 4.224 | 1.364 | 1.614 | 0.929 |

| | | | | | | | | | | |
|---|---|---|---|---|---|---|---|---|---|---|
| Solyc03g117590.3 | Chaperone protein DnaJ (AHRD V3.3 *** A0A2G2WQF7_CAPBA) | NA | -1.613 | 9.79E-08 | 32.380 | 27.355 | 38.166 | 10.478 | 14.769 | 7.260 |
| Solyc06g005140.3 | Pentatricopeptide repeat-containing protein, mitochondrial (AHRD V3.3 *** A0A2G3DBW2_CAPCH) | NA | -1.066 | 1.44E-07 | 10.363 | 10.476 | 9.141 | 4.697 | 4.734 | 4.949 |
| Solyc04g077970.4 | Adenine phosphoribosyltransferase 3 (AHRD V3.3 *** A0A2G3CPQ3_CAPCH) | APRT | -1.074 | 1.64E-07 | 506.273 | 442.197 | 577.031 | 251.806 | 244.071 | 232.352 |
| Solyc01g111120.4 | Triosephosphate isomerase (AHRD V3.3 *** A0A200Q3A8_9MAGN) | TPI | -0.967 | 1.67E-07 | 142.493 | 152.250 | 141.099 | 77.928 | 73.162 | 73.280 |
| Solyc11g072860.2 | Histone H4 (AHRD V3.3 *-* F2E7L1_HORVV) | H4 | -5.843 | 2.01E-07 | 28.029 | 25.509 | 38.786 | 1.078 | 0.479 | 0.000 |
| Solyc02g070800.2 | DUF561 domain-containing protein (AHRD V3.3 *** A0A1Q3DH97_CEPFO) | NA | -2.581 | 2.08E-07 | 19.717 | 10.469 | 29.530 | 4.337 | 3.732 | 2.011 |
| Solyc01g108660.4 | N-acetyl-gamma-glutamyl-phosphate reductase (AHRD V3.3 *** A0A1U8E8L1_CAPAN) | ARGC | -1.016 | 2.14E-07 | 47.047 | 51.229 | 46.647 | 27.642 | 25.630 | 19.297 |
| Solyc05g012510.3 | Alpha-1,4 glucan phosphorylase (AHRD V3.3 *** A0A2G2VL62_CAPBA) | PYG | -2.319 | 2.47E-07 | 9.205 | 5.932 | 10.754 | 2.186 | 2.249 | 0.844 |
| Solyc11g020610.3 | Neutral/alkaline invertase (AHRD V3.3 *** I0CL58_MANES) | NA | -1.570 | 2.94E-07 | 5.004 | 2.951 | 3.527 | 1.248 | 1.375 | 1.243 |
| Solyc07g056500.4 | glutathione S-transferase T5 | GST | -1.204 | 3.13E-07 | 60.210 | 54.075 | 57.020 | 29.389 | 28.634 | 17.407 |
| Solyc09g011630.3 | Glutation-S-transferase | GST | -2.714 | 3.25E-07 | 20.023 | 22.234 | 13.681 | 3.787 | 3.833 | 1.117 |
| Solyc04g018055.1 | Glyceraldehyde-3-phosphate dehydrogenase (AHRD V3.3 *-* A0A2G3C9S8_CAPCH) | GAPDH | -2.467 | 3.43E-07 | 4.096 | 6.072 | 7.185 | 0.620 | 1.145 | 1.352 |
| Solyc09g074880.4 | Protein LOW PSII ACCUMULATION 1, chloroplastic (AHRD V3.3 *** A0A2G2W035_CAPBA) | NA | -1.227 | 3.61E-07 | 6.033 | 6.974 | 6.500 | 2.598 | 3.308 | 2.506 |
| Solyc06g005360.3 | Actin-depolymerizing factor (AHRD V3.3 *** D9I9X9_HEVBR) | CFL | -0.978 | 4.07E-07 | 778.529 | 769.960 | 866.271 | 424.670 | 427.915 | 382.927 |
| Solyc03g026110.4 | SUN-like protein 8 | NA | -1.926 | 4.78E-07 | 11.305 | 9.262 | 14.995 | 4.393 | 3.258 | 1.843 |
| Solyc09g031970.3 | Alpha-1,4 glucan phosphorylase (AHRD V3.3 *** A0A1U8E418_CAPAN) | PYG | -0.992 | 7.14E-07 | 16.948 | 12.349 | 15.489 | 7.523 | 7.331 | 7.697 |
| Solyc01g099190.4 | lipoxygenase B | LOX1_5 | -1.293 | 9.31E-07 | 3201.843 | 3623.665 | 4883.220 | 1723.368 | 1630.951 | 1462.503 |

| | | | | | | | | | | |
|---|---|---|---|---|---|---|---|---|---|---|
| Solyc06g005390.1 | Histone H2B (AHRD V3.3 *** A0A1U8GXA4_CAPAN) | H2B | -1.331 | 9.84E-07 | 18.426 | 17.196 | 17.127 | 6.396 | 6.468 | 8.078 |
| Solyc03g121610.3 | Serine/threonine-protein kinase PBS1 (AHRD V3.3 *** A0A2G2XXF9_CAPAN) | NA | -1.840 | 1.46E-06 | 2.825 | 1.849 | 1.863 | 0.669 | 0.805 | 0.379 |
| Solyc09g005230.3 | Protein root UVB sensitive 5 (AHRD V3.3 *** A0A2G3BL77_CAPCH) | NA | -1.349 | 1.78E-06 | 4.045 | 4.807 | 5.064 | 1.556 | 1.829 | 2.083 |
| Solyc06g008220.4 | Multiple organellar RNA editing factor 2, chloroplastic (AHRD V3.3 *** A0A2G3C3Y9_CAPCH) | NA | -1.269 | 2.07E-06 | 25.438 | 27.357 | 32.458 | 13.683 | 14.046 | 8.189 |
| Solyc07g056490.4 | Glutathione S-transferase-like protein (AHRD V3.3 *** A0A2K3N2T5_TRIPR) | GST | -1.038 | 2.44E-06 | 73.439 | 65.703 | 87.171 | 40.229 | 42.710 | 28.615 |
| Solyc08g080940.3 | glutathione peroxidase like encoding 1 | GPX | -0.943 | 2.81E-06 | 592.349 | 589.887 | 647.342 | 297.302 | 340.219 | 319.930 |
| Solyc01g109040.4 | Cytochrome b6-f complex subunit 7 (AHRD V3.3 *** A0A1U8DXD4_CAPAN) | NA | -1.806 | 2.99E-06 | 6.408 | 6.453 | 4.365 | 1.450 | 1.444 | 1.991 |
| Solyc06g064550.3 | Aspartokinase-homoserine dehydrogenase (AHRD V3.3 *** O63067_SOYBN) | THRA | -1.660 | 3.22E-06 | 3.700 | 3.699 | 2.771 | 1.280 | 1.343 | 0.652 |
| Solyc01g005560.3 | Isocitrate dehydrogenase [NADP] (AHRD V3.3 *** A0A2G3BR90_CAPCH) | IDH1 | -1.386 | 3.28E-06 | 231.684 | 360.484 | 228.468 | 107.432 | 98.964 | 110.199 |
| Solyc03g033330.3 | RING/U-box superfamily protein (AHRD V3.3 *** A0A2U1ML59_ARTAN) | NA | -2.908 | 4.90E-06 | 1.810 | 5.464 | 6.923 | 0.664 | 0.772 | 0.473 |
| Solyc03g093360.3 | PLAT/LH2 domain (AHRD V3.3 *** A0A200QDT2_9MAGN) | NA | -3.679 | 8.29E-06 | 11.956 | 23.698 | 42.717 | 2.911 | 2.816 | 0.573 |
| Solyc09g064800.3 | Isoamylase 2, chloroplastic (AHRD V3.3 *** A0A2G3A6V3_CAPAN) | ISA | -1.028 | 1.06E-05 | 15.028 | 13.303 | 11.659 | 6.111 | 6.080 | 7.422 |
| Solyc10g078920.3 | Thioredoxin-like 3-1, chloroplastic (AHRD V3.3 *** A0A2G3BHN6_CAPCH) | NA | -1.900 | 1.09E-05 | 24.200 | 10.805 | 29.463 | 5.926 | 6.307 | 5.074 |
| Solyc05g016330.3 | Cytochrome P450 (AHRD V3.3 *** A0A200QCI8_9MAGN) | CYP97B3 | -1.049 | 1.22E-05 | 16.535 | 15.672 | 21.278 | 10.113 | 9.290 | 6.734 |
| Solyc03g082580.3 | 6-phosphogluconolactonase (AHRD V3.3 *** B6UAK0_MAIZE) | PGLS | -0.773 | 1.41E-05 | 30.391 | 27.955 | 29.071 | 18.743 | 18.257 | 14.643 |
| Solyc04g015040.3 | Peptidylprolyl isomerase (AHRD V3.3 *** A0A2G2XYE0_CAPAN) | peptidylprolyl isomerase | -1.020 | 1.42E-05 | 12.020 | 13.289 | 13.754 | 6.025 | 6.476 | 6.845 |
| Solyc09g059030.4 | Quinone-oxidoreductase chloroplastic-like (AHRD V3.3 *** A0A2K3P901_TRIPR) | CEQORH | -1.058 | 1.45E-05 | 109.934 | 92.491 | 127.545 | 47.470 | 50.827 | 60.106 |

| | | | | | | | | | | |
|---|---|---|---|---|---|---|---|---|---|---|
| Solyc05g050970.4 | Transketolase (AHRD V3.3 *** A0A200R9X0_9MAGN) | TKTA | -1.112 | 1.48E-05 | 87.110 | 113.234 | 82.375 | 42.408 | 41.256 | 47.819 |
| Solyc02g092940.3 | Receptor protein kinase-like protein (AHRD V3.3 *** W9RY70_9ROSA) | NA | -1.198 | 1.59E-05 | 4.816 | 3.124 | 3.592 | 1.583 | 1.964 | 1.505 |
| Solyc01g081390.4 | Xylulose 5-phosphate/phosphate translocator, chloroplastic (AHRD V3.3 *** A0A1U8EHP7_CAPAN) | SLC35E1 | -1.042 | 1.72E-05 | 20.845 | 23.180 | 18.622 | 11.888 | 11.400 | 7.579 |
| Solyc04g064480.3 | Unknown protein | NA | -1.292 | 2.93E-05 | 7.846 | 7.286 | 7.140 | 3.240 | 2.943 | 2.926 |
| Solyc03g120450.4 | Aminotransferase (AHRD V3.3 *** A0A200PN75_9MAGN) | ISS1 | -1.619 | 3.33E-05 | 3.000 | 3.083 | 2.566 | 0.858 | 1.377 | 0.615 |
| Solyc02g080810.3 | Aminomethyltransferase (AHRD V3.3 *** A0A2G2VBG4_CAPBA) | GCVT | -0.918 | 3.46E-05 | 33.080 | 40.541 | 30.818 | 17.647 | 18.482 | 19.463 |
| Solyc06g075930.1 | Histone H4 (AHRD V3.3 *-* F2E7L1_HORVV) | H4 | -1.332 | 4.10E-05 | 13.266 | 12.000 | 17.933 | 6.411 | 5.945 | 4.838 |
| Solyc04g081440.3 | beta-fructofuranosidase | NA | -0.830 | 4.23E-05 | 83.265 | 74.331 | 98.244 | 48.208 | 50.785 | 45.784 |
| Solyc07g066600.3 | Phosphoglycerate kinase (AHRD V3.3 *** A0A2G3C3B1_CAPCH) | PGK | -1.042 | 4.30E-05 | 322.385 | 364.264 | 483.516 | 204.784 | 195.744 | 172.871 |
| Solyc10g083880.2 | tonoplast intrinsic protein 1.3 | TIP | -1.884 | 5.03E-05 | 51.527 | 42.241 | 18.212 | 8.428 | 11.098 | 10.907 |
| Solyc06g060260.3 | Stromal ascorbate peroxidase (AHRD V3.3 *** Q9TNL9_TOBAC) | L-ascorbate peroxidase | -1.051 | 5.06E-05 | 137.888 | 186.156 | 133.050 | 75.441 | 69.291 | 77.262 |
| Solyc04g054980.3 | PLAT/LH2 domain (AHRD V3.3 *** A0A200QB70_9MAGN) | NA | -1.767 | 5.51E-05 | 62.728 | 72.625 | 92.397 | 31.231 | 26.299 | 10.881 |
| Solyc10g078740.2 | Enoyl-[acyl-carrier-protein] reductase [NADH] (AHRD V3.3 *** B6TFF6_MAIZE) | FABI | -2.833 | 5.82E-05 | 2.651 | 1.783 | 4.431 | 0.552 | 0.576 | 0.126 |
| Solyc09g009020.3 | enolase | ENO | -1.195 | 6.09E-05 | 360.150 | 414.660 | 525.185 | 229.918 | 209.447 | 136.912 |
| Solyc09g083120.3 | Acylamino-acid-releasing enzyme (AHRD V3.3 *** A0A1J3FLE9_NOCCA) | APEH | -1.076 | 6.26E-05 | 2.833 | 2.392 | 2.859 | 1.282 | 1.544 | 1.043 |
| Solyc02g077090.4 | Protein CHUP1, chloroplastic (AHRD V3.3 *-* A0A2G3D559_CAPCH) | NA | -2.191 | 6.72E-05 | 1.759 | 1.718 | 2.066 | 0.616 | 0.456 | 0.155 |
| Solyc01g067750.3 | dehydroquinate dehydratase/shikimate:NADP oxidoreductase | ARODE | -1.129 | 8.02E-05 | 38.142 | 47.184 | 27.008 | 17.611 | 16.719 | 17.435 |
| Solyc03g119970.3 | ADP,ATP carrier protein (AHRD V3.3 *** A0A2G3D148_CAPCH) | TC.AAA | -1.308 | 9.55E-05 | 4.699 | 2.968 | 4.452 | 1.750 | 2.026 | 1.167 |
| Solyc07g015860.3 | peptide deformylase AF271258 | PDF | -1.407 | 1.03E-04 | 2.978 | 3.756 | 4.797 | 1.217 | 1.472 | 1.665 |

| | | | | | | | | | | |
|---|---|---|---|---|---|---|---|---|---|---|
| Solyc12g014180.2 | Malate dehydrogenase (AHRD V3.3 *** A0A2G2X2U9_CAPBA) | MDH2 | -1.180 | 1.10E-04 | 4.934 | 3.771 | 3.495 | 1.645 | 2.149 | 1.621 |
| Solyc01g008160.4 | Protein SLOW GREEN 1, chloroplastic (AHRD V3.3 *** A0A2G2YVV0_CAPAN) | NA | -1.132 | 1.26E-04 | 43.161 | 62.638 | 37.614 | 21.427 | 20.871 | 23.589 |
| Solyc01g111510.3 | Ascorbate peroxidase (AHRD V3.3 *** Q8W4V7_CAPAN) | L-ascorbate peroxidase | -0.697 | 1.26E-04 | 116.017 | 105.661 | 120.658 | 74.170 | 73.879 | 64.650 |
| Solyc01g067730.3 | Acyl carrier protein (AHRD V3.3 *** A0A1U8EET6_CAPAN) | NA | -0.790 | 1.28E-04 | 277.842 | 292.713 | 294.509 | 187.143 | 180.868 | 138.267 |
| Solyc03g116870.3 | NAD(P)H dehydrogenase subunit 48 (AHRD V3.3 *** A0A0F7GZU3_9ROSI) | NA | -1.020 | 1.29E-04 | 8.927 | 9.592 | 12.692 | 5.743 | 5.641 | 4.177 |
| Solyc02g086740.1 | 50S ribosomal protein L12, chloroplastic (AHRD V3.3 *** A0A2G3D9E1_CAPCH) | RP-L7 | -1.037 | 1.59E-04 | 16.258 | 15.242 | 21.160 | 9.734 | 8.977 | 7.154 |
| Solyc06g005230.3 | Receptor-like protein kinase THESEUS 1 (AHRD V3.3 *** A0A2G2WHH2_CAPBA) | NA | -0.878 | 1.67E-04 | 18.611 | 21.581 | 22.635 | 12.826 | 12.783 | 9.046 |
| Solyc08g005570.4 | Root UVB sensitive family (AHRD V3.3 *** A0A2U1QLM3_ARTAN) | NA | -0.910 | 1.69E-04 | 6.094 | 5.952 | 4.695 | 3.081 | 2.939 | 2.939 |
| Solyc03g019880.3 | UPF0426 protein, chloroplastic (AHRD V3.3 *** A0A2G3AW71_CAPCH) | NA | -0.900 | 1.88E-04 | 68.919 | 74.528 | 101.387 | 46.831 | 41.827 | 43.238 |
| Solyc06g051400.3 | omega-3 fatty acid desaturase | FAD3 | -1.214 | 1.92E-04 | 22.364 | 38.346 | 23.258 | 14.221 | 11.568 | 10.880 |
| Solyc03g065340.3 | Alpha-1,4 glucan phosphorylase (AHRD V3.3 *** A0A2G3CMG9_CAPCH) | PYG | -1.376 | 1.95E-04 | 27.997 | 18.000 | 31.672 | 12.435 | 11.799 | 6.129 |
| Solyc06g071290.3 | Aldehyde dehydrogenase (AHRD V3.3 *** A0A0K9R405_SPIOL) | BETB | -2.017 | 2.28E-04 | 1.414 | 3.119 | 2.011 | 0.811 | 0.477 | 0.354 |
| Solyc06g084260.3 | Fatty-acid-binding protein 1 (AHRD V3.3 *** A0A1U8HAC4_CAPAN) | NA | -0.911 | 2.45E-04 | 7.241 | 7.979 | 7.542 | 4.581 | 4.338 | 3.316 |
| Solyc07g020860.3 | Peroxiredoxin (AHRD V3.3 *** Q8S3L0_POPPZ) | PRXII | -0.712 | 2.60E-04 | 319.863 | 338.240 | 337.589 | 208.978 | 213.096 | 190.899 |
| Solyc10g079260.2 | Protein FATTY ACID EXPORT 1, chloroplastic (AHRD V3.3 *** A0A1U8E9B3_CAPAN) | NA | -0.657 | 2.60E-04 | 38.436 | 39.595 | 36.764 | 26.169 | 25.312 | 21.969 |
| Solyc06g073090.3 | chloroplast-specific ribosomal protein chloroplast-specific ribosomal protein | NA | -0.883 | 2.60E-04 | 33.447 | 25.108 | 34.711 | 16.131 | 15.601 | 18.773 |
| Solyc12g099930.2 | Serine--glyoxylate aminotransferase (AHRD V3.3 *** A0A2G2YAX9_CAPAN) | AGXT | -1.108 | 2.60E-04 | 11.734 | 10.534 | 7.205 | 5.290 | 4.311 | 4.151 |
| Solyc04g011510.4 | Triosephosphate isomerase (AHRD V3.3 *** S8E0E9_9LAMI) | TPI | -0.898 | 2.62E-04 | 168.174 | 189.623 | 248.155 | 114.434 | 117.053 | 96.750 |

| | | | | | | | | | | |
|---|---|---|---|---|---|---|---|---|---|---|
| Solyc05g012110.4 | 6-phosphogluconolactonase (AHRD V3.3 *** B6U0H2_MAIZE) | PGLS | -0.774 | 2.93E-04 | 17.786 | 20.337 | 18.917 | 11.321 | 10.557 | 11.624 |
| Solyc06g051410.4 | N-(5'-phosphoribosyl)anthranilate isomerase (AHRD V3.3 *** S8D1B0_9LAMI) | TRPF | -1.675 | 2.95E-04 | 9.994 | 7.367 | 17.754 | 3.912 | 4.756 | 2.469 |
| Solyc01g005800.3 | Calmodulin binding protein (AHRD V3.3 *** A0A2K3N563_TRIPR) | NA | -1.594 | 3.16E-04 | 4.175 | 3.274 | 2.265 | 1.288 | 1.354 | 0.624 |
| Solyc07g006730.4 | Protein DETOXIFICATION (AHRD V3.3 *** A0A2I4FHR8_9ROSI) | TC.MATE | -1.695 | 3.37E-04 | 3.788 | 4.630 | 4.686 | 1.631 | 1.875 | 0.630 |
| Solyc05g010260.4 | 6-phosphogluconate dehydrogenase, decarboxylating (AHRD V3.3 *** A0A1U8F1N4_CAPAN) | PGD | -0.932 | 3.86E-04 | 31.482 | 39.929 | 25.682 | 16.686 | 17.389 | 17.219 |
| Solyc08g065220.3 | glycine decarboxylase p-protein | GLDC | -0.829 | 3.86E-04 | 35.966 | 46.408 | 37.902 | 23.483 | 21.334 | 23.302 |
| Solyc12g089210.2 | Ornithine carbamoyltransferase (AHRD V3.3 *** A0A2G2YA09_CAPAN) | OTC | -1.743 | 4.68E-04 | 19.388 | 29.088 | 30.791 | 6.562 | 4.881 | 11.974 |
| Solyc06g007760.4 | Ycf54-like protein (AHRD V3.3 *** B4FED9_MAIZE) | NA | -1.021 | 5.03E-04 | 28.635 | 33.907 | 43.591 | 21.581 | 15.261 | 15.846 |
| Solyc09g013150.4 | Phosphate transporter (AHRD V3.3 *** A0A2H4H6T2_MALDO) | SLC17A | -1.617 | 5.33E-04 | 243.107 | 146.005 | 97.247 | 62.663 | 64.540 | 33.596 |
| Solyc10g080970.2 | Bifunctional protein FolD 2 (AHRD V3.3 *** A0A1U8EB53_CAPAN) | NA | -1.358 | 5.87E-04 | 20.122 | 14.944 | 20.212 | 9.114 | 8.980 | 3.860 |
| Solyc08g061630.3 | YGGT family protein (AHRD V3.3 *** F8WLD3_CITUN) | YGGT | -1.111 | 5.93E-04 | 2.420 | 2.496 | 2.926 | 1.156 | 1.218 | 1.260 |
| Solyc09g018790.3 | Succinic semialdehyde reductase isofom1 | GLYR | -0.678 | 6.11E-04 | 83.077 | 92.124 | 82.087 | 55.626 | 51.841 | 54.241 |
| Solyc07g042250.3 | chaperonin 21 precursor | GROES | -0.793 | 6.26E-04 | 616.998 | 681.479 | 659.859 | 364.816 | 362.952 | 406.268 |
| Solyc01g108020.3 | Thioredoxin M3, chloroplastic (AHRD V3.3 *** A0A1U8E6Y3_CAPAN) | TRXA | -0.780 | 6.63E-04 | 22.049 | 26.718 | 30.742 | 16.001 | 16.058 | 14.613 |
| Solyc01g108910.4 | maternal effect embryo arrest 14 (AHRD V3.3 *** AT2G15890.1) | NA | -2.459 | 6.83E-04 | 92.058 | 25.661 | 152.517 | 17.003 | 21.359 | 11.208 |
| Solyc05g008800.4 | lipid phosphate phosphatase 2-like (AHRD V3.3 *** A0A1S4AJT8_TOBAC) | DPP1 | -0.758 | 7.56E-04 | 8.654 | 8.710 | 8.832 | 5.508 | 5.679 | 4.449 |
| Solyc02g082260.3 | 3-hydroxy-3-methylglutaryl CoA reductase | HMGCR | -1.150 | 7.63E-04 | 12.820 | 17.088 | 21.533 | 10.076 | 7.903 | 5.555 |
| Solyc02g093830.4 | Glucose-6-phosphate 1-dehydrogenase (AHRD V3.3 *** F2Z9R1_NICBE) | G6PD | -0.671 | 7.69E-04 | 100.435 | 97.712 | 84.832 | 62.006 | 60.205 | 56.745 |

| | | | | | | | | | | |
|---|---|---|---|---|---|---|---|---|---|---|
| Solyc12g014390.3 | 50S ribosomal protein L13 (AHRD V3.3 *** A0A1U8E5A8_CAPAN) | RP-L13 | -0.801 | 7.77E-04 | 46.918 | 47.857 | 66.865 | 31.907 | 30.663 | 30.664 |
| Solyc11g066660.3 | Magnesium transporter MRS2-A, chloroplastic (AHRD V3.3 *** A0A2G2YHE9_CAPAN) | MRS2 | -1.521 | 7.95E-04 | 5.538 | 14.023 | 9.124 | 3.529 | 2.987 | 3.576 |
| Solyc05g053810.3 | Serine hydroxymethyltransferase (AHRD V3.3 *** A0A2G3CI35_CAPCH) | GLYA | -0.873 | 7.99E-04 | 54.081 | 59.668 | 45.602 | 33.682 | 32.812 | 21.855 |
| Solyc08g077530.3 | Beta-amylase (AHRD V3.3 *** Q94EU9_SOLTU) | beta-amylase | -0.920 | 8.28E-04 | 50.661 | 31.349 | 43.467 | 18.687 | 26.924 | 21.012 |
| Solyc01g105020.3 | Protein phosphatase 2C (PP2C)-like domain (AHRD V3.3 *-* A0A200PXZ2_9MAGN) | NA | -1.154 | 9.30E-04 | 1.615 | 1.428 | 2.149 | 0.897 | 0.904 | 0.561 |
| Solyc12g005180.2 | Chloroplast lipocalin (AHRD V3.3 *** Q38JB4_SOLTU) | NA | -0.727 | 9.65E-04 | 31.490 | 22.983 | 32.127 | 17.273 | 18.130 | 17.071 |
| Solyc11g009080.2 | DAHP synthase 1 precursor | AROF | -0.836 | 1.17E-03 | 119.457 | 91.230 | 111.477 | 54.913 | 55.857 | 69.300 |
| Solyc03g113780.3 | NAD(P)H-hydrate epimerase (AHRD V3.3 *** A0A2G2VFG5_CAPBA) | PPOX | -0.594 | 1.21E-03 | 16.881 | 17.737 | 18.332 | 12.139 | 12.251 | 10.979 |
| Solyc07g063190.3 | Thioredoxin (AHRD V3.3 *** A0A200PQX5_9MAGN) | TRXA | -0.704 | 1.30E-03 | 91.617 | 99.769 | 115.400 | 63.000 | 59.589 | 66.372 |
| Solyc05g005180.3 | 1,4-dihydroxy-2-naphthoyl-CoA synthase (AHRD V3.3 *** A0A2G2XZJ4_CAPAN) | MENB | -1.099 | 1.33E-03 | 4.050 | 3.498 | 2.674 | 1.369 | 1.729 | 1.683 |
| Solyc06g074530.1 | Arogenate dehydratase (AHRD V3.3 *** A0A2G2VAY6_CAPBA) | ADT | -2.391 | 1.34E-03 | 320.384 | 210.375 | 166.554 | 58.261 | 67.023 | 11.498 |
| Solyc11g006910.3 | Ferredoxin (AHRD V3.3 *** K4D4V2_SOLLC) | PETF | -0.794 | 1.34E-03 | 45.010 | 36.826 | 33.549 | 22.775 | 21.401 | 22.558 |
| Solyc02g079960.3 | Thioredoxin (AHRD V3.3 *** A0A2G3D6G6_CAPCH) | TRXA | -0.695 | 1.39E-03 | 84.635 | 87.387 | 113.182 | 61.721 | 59.283 | 56.326 |
| Solyc04g071650.4 | Cellulose synthase (AHRD V3.3 *** K4BTF5_SOLLC) | CESA | -1.889 | 1.41E-03 | 4.441 | 4.957 | 4.703 | 1.981 | 1.481 | 0.442 |
| Solyc09g075290.3 | 60S ribosomal protein L18 (AHRD V3.3 *** B6SJC8_MAIZE) | RP-L18E | -0.639 | 1.49E-03 | 130.159 | 139.499 | 153.260 | 88.915 | 94.366 | 90.086 |
| Solyc01g005390.3 | Nudix hydrolase (AHRD V3.3 *** A0A2U1MBD3_ARTAN) | diphosphoinositol-polyphosphate diphosphatase | -2.582 | 1.50E-03 | 57.338 | 23.489 | 19.952 | 7.217 | 8.413 | 1.565 |
| Solyc04g076810.4 | Non-specific serine/threonine protein kinase (AHRD V3.3 *** A0A2G3CPB2_CAPCH) | NA | -0.567 | 1.53E-03 | 21.833 | 22.183 | 23.254 | 15.576 | 16.113 | 14.085 |

| | | | | | | | | | | |
|---|---|---|---|---|---|---|---|---|---|---|
| Solyc03g111010.4 | Glyceraldehyde-3-phosphate dehydrogenase (AHRD V3.3 *** A0A0F7JIC2_NICBE) | GAPDH | -0.695 | 1.60E-03 | 233.911 | 262.767 | 220.247 | 147.305 | 149.398 | 149.118 |
| Solyc03g111000.3 | Glyceraldehyde-3-phosphate dehydrogenase (AHRD V3.3 *** A0A2G2X869_CAPBA) | GAPDH | -0.884 | 1.64E-03 | 6.522 | 4.802 | 6.342 | 3.386 | 3.420 | 2.813 |
| Solyc05g023900.1 | Pentatricopeptide repeat (AHRD V3.3 *** A0A200R3X1_9MAGN) | NA | -0.974 | 1.64E-03 | 1.596 | 1.415 | 1.670 | 0.717 | 0.820 | 0.851 |
| Solyc09g059040.3 | Oxidoreductase, zinc-binding dehydrogenase family protein (AHRD V3.3 *** A0A2U1PT68_ARTAN) | CEQORH | -0.633 | 1.78E-03 | 219.533 | 185.711 | 203.842 | 135.858 | 132.024 | 126.720 |
| Solyc04g076790.3 | Serine hydroxymethyltransferase (AHRD V3.3 *** A0A2G2ZR28_CAPAN) | GLYA | -0.760 | 1.80E-03 | 76.912 | 96.614 | 71.308 | 49.185 | 47.472 | 48.993 |
| Solyc07g056480.3 | glutathione S-transferase/peroxidase | GST | -1.409 | 1.83E-03 | 13.507 | 10.344 | 5.393 | 3.730 | 4.421 | 2.968 |
| Solyc10g083760.3 | chloroplast threonine deaminase 1 precursor | ILVA | -1.026 | 1.96E-03 | 27.340 | 28.081 | 48.111 | 18.706 | 19.251 | 13.481 |
| Solyc07g063000.4 | Phytosulfokine receptor 2 (AHRD V3.3 *** A0A2G3C1N9_CAPCH) | NA | -1.039 | 2.06E-03 | 1.941 | 1.397 | 2.385 | 1.054 | 0.939 | 0.804 |
| Solyc04g005160.3 | 6-phosphogluconate dehydrogenase, decarboxylating (AHRD V3.3 *** A0A1U8F1N4_CAPAN) | PGD | -1.176 | 2.35E-03 | 207.363 | 185.132 | 135.588 | 97.118 | 95.115 | 45.906 |
| Solyc10g081510.2 | ethylene-responsive methionine synthase | METE | -0.852 | 2.37E-03 | 166.531 | 233.780 | 158.282 | 107.507 | 110.213 | 95.400 |
| Solyc02g082760.3 | ethylene-responsive catalase | KATE | -0.972 | 2.37E-03 | 21.519 | 18.217 | 14.153 | 8.571 | 7.660 | 11.146 |
| Solyc11g068430.3 | Ferredoxin (AHRD V3.3 *** K4DA01_SOLLC) | PETF | -0.987 | 2.48E-03 | 527.000 | 546.083 | 926.256 | 330.849 | 378.507 | 307.692 |
| Solyc10g005180.3 | Ycf3-interacting protein 1, chloroplastic (AHRD V3.3 *** Y3IP1_TOBAC) | NA | -0.918 | 2.52E-03 | 40.569 | 41.844 | 50.999 | 29.545 | 26.625 | 15.579 |
| Solyc03g005330.1 | Non-specific serine/threonine protein kinase (AHRD V3.3 *** A0A2G2WWB5_CAPBA) | SNF1 | -0.979 | 2.61E-03 | 3.947 | 4.046 | 5.671 | 2.570 | 2.496 | 1.925 |
| Solyc02g065380.3 | Cold-regulated plasma membrane protein 2 (AHRD V3.3 *** A0A1U8FJM9_CAPAN) | NA | -0.539 | 2.64E-03 | 57.524 | 54.005 | 54.970 | 41.553 | 38.596 | 35.242 |
| Solyc06g062840.4 | Threonine synthase, chloroplastic (AHRD V3.3 *** A0A1U8H4Z2_CAPAN) | THRC | -1.025 | 2.65E-03 | 4.760 | 5.046 | 4.231 | 2.936 | 2.558 | 1.519 |

| | | | | | | | | | | |
|---|---|---|---|---|---|---|---|---|---|---|
| Solyc09g009390.3 | Monodehydroascorbate reductase (AHRD V3.3 *** A0A2G2VWV8_CAPBA) | Monodehydro ascorbate reductase (NADH) | -0.617 | 2.78E-03 | 231.370 | 226.525 | 223.584 | 149.085 | 148.469 | 149.104 |
| Solyc07g062030.3 | Chalcone-flavonone isomerase family protein (AHRD V3.3 *** A0A2G2WEX9_CAPBA) | NA | -0.786 | 2.83E-03 | 9.430 | 10.529 | 13.193 | 6.795 | 6.494 | 6.084 |
| Solyc09g009940.3 | Signal recognition particle protein (AHRD V3.3 *** A0A1U8EGH6_CAPAN) | SRP54 | -0.545 | 2.84E-03 | 41.808 | 38.626 | 43.044 | 28.669 | 28.427 | 27.943 |
| Solyc06g009020.2 | Glutathione S-transferase (AHRD V3.3 *** A0A2G3C3Q3_CAPCH) | GST | -0.945 | 2.93E-03 | 605.668 | 852.794 | 579.066 | 325.559 | 335.066 | 402.223 |
| Solyc10g017850.3 | Peroxisomal membrane protein 11C (AHRD V3.3 *** A0A2G3BDG9_CAPCH) | NA | -0.694 | 3.59E-03 | 10.954 | 12.152 | 10.170 | 7.412 | 6.946 | 6.382 |
| Solyc04g009530.4 | Glutathione S-transferase (AHRD V3.3 *** A0A200QHH9_9MAGN) | GST | -1.075 | 3.60E-03 | 2.488 | 3.192 | 3.866 | 1.732 | 1.612 | 1.231 |
| Solyc06g074510.4 | Phosphoglycerate/bisphosphoglycerate mutase family protein (AHRD V3.3 *** A0A1P8BB43_ARATH) | 2-carboxy-D-arabinitol-1-phosphatase | -0.882 | 4.41E-03 | 2.813 | 3.236 | 3.229 | 1.682 | 2.081 | 1.332 |
| Solyc12g010040.2 | leucine aminopeptidase A | CARP | -0.589 | 4.45E-03 | 195.360 | 198.213 | 202.384 | 138.542 | 132.813 | 127.520 |
| Solyc06g082750.3 | 50S ribosomal protein L17 (AHRD V3.3 *** A0A1U8H2A3_CAPAN) | RP-L17 | -0.754 | 4.64E-03 | 47.452 | 43.344 | 67.716 | 29.661 | 34.319 | 30.545 |
| Solyc11g005620.1 | Acetylglutamate kinase (AHRD V3.3 *** A0A1U8EW67_CAPAN) | ARGB | -0.810 | 4.83E-03 | 14.595 | 19.545 | 14.023 | 9.840 | 8.488 | 9.354 |
| Solyc07g007310.3 | Methyltransferase-like (AHRD V3.3 *** Q67W64_ORYSJ) | NA | -0.626 | 5.32E-03 | 17.350 | 20.060 | 19.959 | 13.083 | 11.624 | 12.654 |
| Solyc06g048730.3 | Uroporphyrinogen decarboxylase (AHRD V3.3 *** A0A2G3CAB7_CAPCH) | HEME | -0.624 | 5.35E-03 | 36.791 | 43.479 | 36.081 | 24.430 | 24.671 | 26.777 |
| Solyc02g089750.4 | Proline-rich receptor-like protein kinase PERK9 (AHRD V3.3 *-* A0A2G3D983_CAPCH) | NA | -1.159 | 5.43E-03 | 9.368 | 9.042 | 7.367 | 4.410 | 5.024 | 2.303 |
| Solyc11g071280.2 | 4-amino-4-deoxychorismate lyase | ADCL | -0.795 | 5.66E-03 | 30.181 | 33.861 | 45.666 | 24.603 | 23.096 | 16.354 |
| Solyc08g065480.3 | Ferrochelatase (AHRD V3.3 *** A0A2G3BRU7_CAPCH) | HEMH | -0.641 | 5.81E-03 | 37.146 | 29.318 | 25.346 | 20.537 | 20.643 | 18.069 |
| Solyc10g085030.1 | Heme-binding-like protein (AHRD V3.3 *** A0A2I0X8X3_9ASPA) | NA | -1.104 | 5.86E-03 | 43.526 | 36.065 | 31.874 | 23.107 | 19.991 | 9.675 |

| | | | | | | | | | | |
|---|---|---|---|---|---|---|---|---|---|---|
| Solyc04g009200.3 | glutamate 1-semialdehyde 2,1-aminomutase | HEML | -0.495 | 5.88E-03 | 44.326 | 42.294 | 43.723 | 30.929 | 32.141 | 29.999 |
| Solyc10g039270.2 | Peptidylprolyl isomerase (AHRD V3.3 *** A0A1U8FL13_CAPAN) | NA | -0.671 | 6.09E-03 | 8.462 | 9.350 | 9.496 | 5.734 | 5.831 | 5.688 |
| Solyc06g074240.3 | Beta-carotene,Pfam:PF05834 | CCS1 | -1.107 | 6.31E-03 | 11.689 | 13.105 | 10.407 | 7.144 | 6.508 | 3.030 |
| Solyc10g051373.1 | RNA-binding protein (AHRD V3.3 *-* O24106_NICGU) | CIRBP | -1.059 | 6.32E-03 | 61.442 | 34.226 | 64.887 | 22.176 | 35.387 | 20.245 |
| Solyc12g095850.2 | Tryptophan--tRNA ligase (AHRD V3.3 *** A0A2G2VHY5_CAPBA) | WARS | -1.070 | 6.48E-03 | 1.930 | 1.438 | 2.130 | 1.071 | 0.765 | 0.778 |
| Solyc01g088600.4 | protein TRIGALACTOSYLDIACYLGLYCEROL 4, chloroplastic (AHRD V3.3 *** A0A2I4EI43_9ROSI) | NA | -1.411 | 6.65E-03 | 2.469 | 2.140 | 4.210 | 1.487 | 1.220 | 0.649 |
| Solyc08g081570.3 | 2-C-methyl-D-erythritol 2,4-cyclodiphosphate synthase (AHRD V3.3 *** A0A2G3DAU7_CAPCH) | ISPF | -0.521 | 7.13E-03 | 36.499 | 31.791 | 31.434 | 24.490 | 24.470 | 21.056 |
| Solyc02g085760.2 | Rhomboid domain-containing protein (AHRD V3.3 *** A0A1Q3C608_CEPFO) | NA | -1.250 | 7.16E-03 | 5.459 | 3.246 | 6.360 | 2.880 | 2.164 | 1.351 |
| Solyc01g102820.4 | 2-C-methyl-D-erythritol 4-phosphate cytidylyltransferase (AHRD V3.3 *** A9ZN09_HEVBR) | ISPD | -0.595 | 7.22E-03 | 16.732 | 13.613 | 13.852 | 10.342 | 9.824 | 9.227 |
| Solyc04g073990.3 | annexin p34 | ANNAT | -1.012 | 7.31E-03 | 251.542 | 251.529 | 312.091 | 167.136 | 166.375 | 78.535 |
| Solyc08g077050.3 | Ferredoxin (AHRD V3.3 *** A0A1U8GXA1_CAPAN) | PETF | -0.647 | 7.66E-03 | 61.529 | 55.838 | 58.593 | 32.721 | 36.623 | 43.033 |
| Solyc04g076380.4 | NADPH--cytochrome P450 reductase (AHRD V3.3 *** A0A2G3CP38_CAPCH) | POR | -0.579 | 7.95E-03 | 49.083 | 39.319 | 52.732 | 32.832 | 33.184 | 29.049 |
| Solyc02g083810.4 | Ferredoxin--NADP reductase, chloroplastic (AHRD V3.3 *** K4BAP9_SOLLC) | PETH | -0.662 | 8.09E-03 | 61.270 | 64.614 | 70.064 | 38.904 | 37.633 | 47.377 |
| Solyc01g006450.4 | Enoyl-[acyl-carrier-protein] reductase [NADH], chloroplastic (AHRD V3.3 *** A0A2G3DE80_CAPCH) | FABI | -0.615 | 8.15E-03 | 42.634 | 49.366 | 52.155 | 30.592 | 29.736 | 34.114 |
| Solyc09g092260.4 | Chaperone protein DnaJ (AHRD V3.3 *** A0A2G3AV72_CAPCH) | NA | -1.656 | 8.17E-03 | 6.641 | 4.557 | 16.113 | 2.704 | 3.528 | 2.482 |
| Solyc07g008530.1 | Tyrosine--tRNA ligase (AHRD V3.3 *** A0A1U8H9E5_CAPAN) | YARS | -0.628 | 8.35E-03 | 13.364 | 13.757 | 17.007 | 9.876 | 9.091 | 9.699 |

| | | | | | | | | | | |
|---|---|---|---|---|---|---|---|---|---|---|
| Solyc08g077880.3 | Chlorophyll A-B binding protein (AHRD V3.3 *** A0A200RAS0_9MAGN) | NA | -0.588 | 8.96E-03 | 41.252 | 45.157 | 51.328 | 31.603 | 28.476 | 31.905 |
| Solyc09g072880.3 | Protein FLUORESCENT IN BLUE LIGHT, chloroplastic (AHRD V3.3 *** A0A1U8GA42_CAPAN) | NA | -0.860 | 9.16E-03 | 1.487 | 1.419 | 1.793 | 0.830 | 0.792 | 0.963 |
| Solyc11g007160.2 | RNA-binding (RRM/RBD/RNP motifs) family protein (AHRD V3.3 *** A0A1I9LPX6_ARATH) | NA | -1.028 | 9.21E-03 | 2.142 | 1.549 | 2.074 | 1.128 | 1.023 | 0.697 |
| Solyc11g040110.2 | Cobalt ion binding (AHRD V3.3 *** B6SJF4_MAIZE) | NA | -0.493 | 9.56E-03 | 53.820 | 53.061 | 60.187 | 39.339 | 41.297 | 38.793 |
| Solyc09g075720.4 | Protein kinase superfamily protein (AHRD V3.3 *** Q0WSF6_ARATH) | NA | -1.582 | 9.68E-03 | 2.033 | 3.353 | 4.733 | 1.471 | 1.423 | 0.549 |
| Solyc06g008510.3 | Cdt1-like protein chloroplastic-like (AHRD V3.3 *** A0A2K3MWV0_TRIPR) | CDT1 | -1.830 | 9.79E-03 | 0.767 | 1.970 | 1.273 | 0.211 | 0.342 | 0.560 |
| Solyc08g081010.3 | gamma-glutamylcysteine synthetase 1 | GSHA | -0.709 | 9.87E-03 | 434.731 | 458.018 | 356.851 | 236.922 | 248.055 | 281.679 |
| Solyc01g087040.2 | Photosystem II PsbP (AHRD V3.3 *** A0A200Q959_9MAGN) | NA | -0.804 | 1.01E-02 | 9.438 | 10.624 | 9.567 | 4.706 | 5.634 | 6.652 |
| Solyc10g008740.3 | Mg-protoporphyrin IX chelatase (AHRD V3.3 *** A0A2G2YJA3_CAPAN) | CHLI | -0.609 | 1.01E-02 | 21.548 | 23.403 | 29.819 | 16.607 | 16.694 | 16.017 |
| Solyc05g014470.3 | glyceraldehyde 3-phosphate dehydrogenase | GAPDH | -0.639 | 1.03E-02 | 556.002 | 586.383 | 653.926 | 426.404 | 427.923 | 313.009 |
| Solyc04g076870.4 | Glutamyl-tRNA reductase (AHRD V3.3 *** A6Q0F0_TOBAC) | HEMA | -0.595 | 1.08E-02 | 32.064 | 34.404 | 36.400 | 21.766 | 21.077 | 25.370 |
| Solyc09g082970.3 | Pyruvate kinase (AHRD V3.3 *** A0A2G2W137_CAPBA) | PK | -0.612 | 1.17E-02 | 75.084 | 76.969 | 94.390 | 54.885 | 50.811 | 56.069 |
| Solyc04g053130.4 | Stress enhanced protein 2, chloroplastic (AHRD V3.3 *** A0A1U8F0B2_CAPAN) | NA | -1.101 | 1.17E-02 | 559.495 | 651.886 | 568.270 | 213.388 | 208.379 | 400.834 |
| Solyc08g048550.3 | Protease Do-like 5, chloroplastic (AHRD V3.3 *** A0A1U8DTN0_CAPAN) | NA | -0.532 | 1.21E-02 | 18.079 | 15.048 | 18.203 | 12.284 | 12.199 | 11.219 |
| Solyc10g006310.3 | Protein FATTY ACID EXPORT 6 (AHRD V3.3 *** A0A1U8EPM9_CAPAN) | NA | -0.779 | 1.24E-02 | 9.296 | 9.439 | 9.862 | 6.186 | 6.704 | 4.013 |
| Solyc01g028810.3 | Beta chaperonin 60 (AHRD V3.3 *** Q0W9E2_SOLCO) | GROEL | -0.630 | 1.32E-02 | 661.899 | 721.340 | 680.052 | 434.201 | 421.567 | 481.636 |

| | | | | | | | | | | |
|---|---|---|---|---|---|---|---|---|---|---|
| Solyc11g010480.2 | Protein CURVATURE THYLAKOID 1A, chloroplastic (AHRD V3.3 *** A0A2G3B0C6_CAPCH) | NA | -0.621 | 1.35E-02 | 30.497 | 32.428 | 28.597 | 17.797 | 18.868 | 22.917 |
| Solyc02g068090.3 | 30S ribosomal protein S21 (AHRD V3.3 *** A0A2G2XI83_CAPBA) | NA | -0.562 | 1.40E-02 | 57.387 | 63.259 | 70.237 | 43.578 | 40.323 | 45.886 |
| Solyc03g083440.4 | Glutamate synthase (AHRD V3.3 *** A0A2U1PKW7_ARTAN) | GLT1 | -0.794 | 1.41E-02 | 3.837 | 5.121 | 5.455 | 2.363 | 3.752 | 2.323 |
| Solyc10g079470.3 | gldhL-galactono-1,4-lactone dehydrogenase | GLDH | -0.825 | 1.43E-02 | 5.614 | 5.320 | 7.498 | 3.522 | 4.512 | 2.526 |
| Solyc06g076510.3 | Phosphoserine phosphatase, chloroplastic (AHRD V3.3 *** A0A2G3CBM8_CAPCH),Pfam:PF00702 | SERB | -0.696 | 1.44E-02 | 6.502 | 8.326 | 7.616 | 4.634 | 5.266 | 4.111 |
| Solyc10g083350.2 | Heme-binding-like protein, chloroplastic (AHRD V3.3 *** A0A1U8EAT6_CAPAN) | NA | -0.683 | 1.44E-02 | 11.903 | 9.821 | 13.288 | 7.640 | 6.677 | 7.524 |
| Solyc06g073260.3 | Chloroplast stem-loop binding protein of 41 kDa b, chloroplastic (AHRD V3.3 *** A0A2G3CAL3_CAPCH) | NA | -1.097 | 1.50E-02 | 1.349 | 1.892 | 1.308 | 0.920 | 0.625 | 0.599 |
| Solyc01g100360.4 | Dihydrolipoyl dehydrogenase (AHRD V3.3 *** A0A2U1LTA9_ARTAN) | DLD | -0.529 | 1.52E-02 | 57.020 | 64.417 | 56.340 | 45.323 | 41.782 | 37.261 |
| Solyc05g009010.1 | Protein kinase family protein (AHRD V3.3 *-* A0A2U1MH78_ARTAN) | NA | -1.306 | 1.52E-02 | 22.320 | 16.943 | 13.709 | 9.699 | 8.989 | 3.190 |
| Solyc10g008560.3 | Protein root UVB sensitive 6 (AHRD V3.3 *** A0A2G2VRX7_CAPBA) | NA | -0.448 | 1.56E-02 | 28.938 | 27.208 | 29.087 | 20.590 | 22.114 | 20.175 |
| Solyc11g008680.2 | Acyl-[acyl-carrier-protein] desaturase (AHRD V3.3 *** A0A2G3B0E0_CAPCH) | FAB2 | -1.045 | 1.58E-02 | 23.709 | 24.899 | 50.997 | 14.963 | 15.704 | 17.656 |
| Solyc08g014130.3 | Isopropylmalate synthase (AHRD V3.3 *** K4CJ46_SOLLC) | LEUA | -1.148 | 1.70E-02 | 743.706 | 895.389 | 653.137 | 267.489 | 248.455 | 508.781 |
| Solyc12g098890.2 | 50S ribosomal protein L18 (AHRD V3.3 *** A0A1U8F639_CAPAN) | RP-L18 | -0.616 | 1.72E-02 | 22.809 | 25.426 | 28.160 | 16.110 | 15.962 | 17.932 |
| Solyc05g008930.3 | Protein kinase family protein (AHRD V3.3 *** A0A1P8BDK9_ARATH) | NA | -0.638 | 1.73E-02 | 3.809 | 3.668 | 4.073 | 2.196 | 2.941 | 2.343 |
| Solyc01g009990.3 | Peptidyl-prolyl cis-trans isomerase (AHRD V3.3 *** A0A2G3DGA1_CAPCH) | PPIB | -0.531 | 1.80E-02 | 126.366 | 133.705 | 121.647 | 87.060 | 85.543 | 92.698 |
| Solyc09g020130.3 | 60S ribosomal protein L5-like (AHRD V3.3 *** A0A2I4G950_9ROSI) | RP-L5E | -0.494 | 1.86E-02 | 94.008 | 105.146 | 101.033 | 74.187 | 71.681 | 68.887 |

| | | | | | | | | | | |
|---|---|---|---|---|---|---|---|---|---|---|
| Solyc12g008470.2 | Cytochrome B5-like protein (AHRD V3.3 *** A0A1U8EAT2_CAPAN) | NA | -1.450 | 1.87E-02 | 12.338 | 11.800 | 30.132 | 7.211 | 9.113 | 3.891 |
| Solyc08g079820.3 | Nudix hydrolase 14, chloroplastic (AHRD V3.3 *** A0A2G3AE90_CAPAN) | NUDX14 | -0.453 | 1.90E-02 | 25.317 | 24.080 | 24.507 | 18.627 | 18.921 | 16.855 |
| Solyc03g112770.3 | Glutaredoxin (AHRD V3.3 *** I2FJT7_SOLTU) | GRXC | -0.545 | 1.97E-02 | 24.497 | 23.422 | 30.176 | 19.216 | 19.082 | 15.687 |
| Solyc07g032740.4 | Aspartate aminotransferase (AHRD V3.3 *** A0A0J8BGE0_BETVU) | GOT1 | -1.079 | 1.98E-02 | 59.367 | 51.147 | 123.837 | 40.607 | 40.680 | 30.651 |
| Solyc03g111850.3 | Indole-3-glycerol phosphate synthase (AHRD V3.3 *** A0A2G3CCR7_CAPCH) | TRPC | -0.438 | 2.02E-02 | 35.663 | 34.091 | 36.135 | 27.505 | 26.188 | 24.974 |
| Solyc10g080320.3 | Adenylosuccinate synthetase (AHRD V3.3 *** S8CIB7_9LAMI) | PURA | -0.482 | 2.05E-02 | 42.268 | 46.228 | 40.192 | 32.568 | 30.554 | 29.696 |
| Solyc08g007225.1 | Phospholipase A1-Ibeta2, chloroplastic (AHRD V3.3 *** A0A2G3BPM7_CAPCH) | NA | -0.886 | 2.09E-02 | 1.221 | 1.525 | 1.608 | 0.827 | 0.745 | 0.785 |
| Solyc03g083580.3 | Peptidase_M3 domain-containing protein (AHRD V3.3 *** A0A1Q3BAD5_CEPFO) | MIPEP | -0.739 | 2.11E-02 | 1.884 | 2.192 | 2.604 | 1.346 | 1.321 | 1.357 |
| Solyc01g105160.4 | Copper-transporting ATPase PAA1, chloroplastic (AHRD V3.3 *** A0A2G2X1Z0_CAPBA) | COPA | -0.570 | 2.16E-02 | 16.447 | 20.535 | 16.057 | 11.587 | 11.729 | 12.625 |
| Solyc01g060070.3 | Outer envelope pore protein 16-2, chloroplastic (AHRD V3.3 *** A0A2G2XR36_CAPBA),Pfam:PF02466 | NA | -1.083 | 2.19E-02 | 2.570 | 1.946 | 3.077 | 1.015 | 1.684 | 0.908 |
| Solyc12g042950.2 | ADP,ATP carrier protein (AHRD V3.3 *** A0A2G2Y8J4_CAPAN) | TC.AAA | -0.533 | 2.20E-02 | 56.730 | 41.984 | 45.784 | 36.206 | 36.336 | 28.181 |
| Solyc05g008860.4 | Receptor protein kinase-like protein (AHRD V3.3 *** A0A2R6P9S7_ACTCH) | NA | -0.737 | 2.20E-02 | 1.511 | 1.683 | 1.646 | 1.060 | 0.879 | 0.978 |
| Solyc07g007600.3 | vacuolar-type H+-pyrophosphatase | AVP | -0.922 | 2.23E-02 | 87.784 | 42.173 | 51.027 | 34.057 | 38.420 | 23.857 |
| Solyc02g086730.3 | Ribosomal protein L7/L12 (AHRD V3.3 *** A0A200R2P7_9MAGN) | RP-L7 | -0.518 | 2.23E-02 | 86.269 | 84.717 | 101.338 | 61.380 | 61.313 | 67.959 |
| Solyc10g006010.3 | Two pore potassium channel a (AHRD V3.3 *** A0A2G2VRA3_CAPBA) | KCNKF | -0.628 | 2.24E-02 | 5.268 | 5.828 | 6.252 | 4.056 | 4.216 | 3.084 |
| Solyc09g015370.1 | Pentatricopeptide repeat (AHRD V3.3 *** A0A200R528_9MAGN) | NA | -1.002 | 2.26E-02 | 1.614 | 1.496 | 1.580 | 0.592 | 0.695 | 1.037 |
| Solyc08g082250.3 | endo-beta-1,4-D-glucanase (Cel8) | endoglucanase | -1.428 | 2.29E-02 | 6.018 | 1.803 | 7.048 | 1.729 | 1.957 | 1.824 |

| | | | | | | | | | | |
|---|---|---|---|---|---|---|---|---|---|---|
| Solyc06g074430.4 | 60S acidic ribosomal protein P2 (AHRD V3.3 *** A0A2G2ZDK5_CAPAN),Pfam:PF00428 | RP-LP2 | -0.492 | 2.37E-02 | 231.070 | 230.116 | 264.280 | 182.120 | 188.110 | 150.789 |
| Solyc08g082820.4 | TOMBIPGRBC Tomato BiP (binding protein)/grp78 | HSPA5 | -0.689 | 2.53E-02 | 33.679 | 55.334 | 40.643 | 27.582 | 27.998 | 25.770 |
| Solyc01g098650.2 | GDSL esterase/lipase (AHRD V3.3 *** A0A2G3BUB9_CAPCH) | NA | -0.580 | 2.59E-02 | 27.391 | 29.400 | 37.714 | 24.522 | 21.108 | 18.173 |
| Solyc02g085130.4 | Tubby-like F-box protein (AHRD V3.3 *** A0A2G3D7T8_CAPCH) | TUB | -1.251 | 2.63E-02 | 0.920 | 1.637 | 1.771 | 0.684 | 0.801 | 0.366 |
| Solyc06g071070.1 | NAD(P)-binding Rossmann-fold superfamily protein (AHRD V3.3 *** A0A2U1L921_ARTAN),Pfam:PF13561 | FABG | -0.865 | 2.63E-02 | 7.913 | 6.306 | 4.830 | 3.285 | 4.392 | 2.885 |
| Solyc12g043040.1 | Sulfate transporter (AHRD V3.3 *** A1YKF8_BRASY) | SULTR3 | -0.865 | 2.65E-02 | 3.425 | 2.473 | 3.451 | 1.727 | 1.419 | 1.958 |
| Solyc01g105560.3 | GTP cyclohydrolase II/3,4-dihydroxy-2-butanone 4-phosphate synthase | RIBBA | -0.554 | 2.74E-02 | 134.488 | 119.049 | 106.411 | 89.148 | 91.754 | 67.063 |
| Solyc03g121720.2 | Glyoxylate reductase 2 (AHRD V3.3 *** A0A1P8AS42_ARATH) | GLYR | -0.969 | 2.79E-02 | 2.523 | 2.166 | 1.543 | 0.869 | 1.114 | 1.203 |
| Solyc10g084770.3 | Protein kinase APK1B, chloroplastic (AHRD V3.3 *** A0A1U8E7T5_CAPAN) | NA | -1.169 | 2.83E-02 | 2.128 | 1.487 | 1.700 | 1.024 | 0.994 | 0.389 |
| Solyc08g066850.3 | Lactoylglutathione lyase (AHRD V3.3 *** A0A1U8DS55_CAPAN) | GLO1 | -0.620 | 2.94E-02 | 2.324 | 2.141 | 2.151 | 1.540 | 1.333 | 1.443 |
| Solyc08g069030.4 | aminolevulinic acid dehydratase | HEMB | -0.522 | 2.95E-02 | 104.334 | 105.612 | 105.316 | 72.653 | 67.987 | 79.425 |
| Solyc04g078850.4 | defective chloroplasts and leaves | NA | -0.481 | 3.15E-02 | 17.702 | 19.521 | 18.046 | 13.487 | 13.161 | 13.201 |
| Solyc01g098610.4 | glutathione synthetase 2 | GSS | -0.485 | 3.18E-02 | 10.502 | 9.045 | 8.538 | 6.949 | 7.155 | 6.105 |
| Solyc09g014400.3 | Phosphoglucan, water dikinase, chloroplastic (AHRD V3.3 *-* A0A1J3J5B0_NOCCA) | NA | -0.634 | 3.18E-02 | 4.943 | 4.876 | 6.375 | 3.491 | 3.426 | 3.560 |
| Solyc10g086190.3 | Adenosine kinase (AHRD V3.3 *** A0A1U8FGB1_CAPAN) | ADK | -1.360 | 3.21E-02 | 4.461 | 6.885 | 4.462 | 3.000 | 2.498 | 0.829 |
| Solyc05g013380.3 | Alanine aminotransferase 2 (AHRD V3.3 *** B6TXZ8_MAIZE) | GGAT | -0.813 | 3.38E-02 | 7.949 | 9.734 | 11.047 | 6.412 | 6.852 | 3.412 |
| Solyc09g011620.2 | Glutathione S-transferase-like protein (AHRD V3.3 *** A8DUB0_SOLLC) | GST | -0.624 | 3.43E-02 | 8.278 | 7.282 | 7.508 | 4.828 | 5.417 | 4.789 |

| | | | | | | | | | | |
|---|---|---|---|---|---|---|---|---|---|---|
| Solyc02g063270.4 | Protein DETOXIFICATION (AHRD V3.3 *** A0A328DLR5_9ASTE) | TC.MATE | -0.682 | 3.45E-02 | 50.153 | 40.415 | 69.635 | 37.025 | 37.497 | 26.352 |
| Solyc01g110550.3 | Ribosome-binding factor A (AHRD V3.3 *** A0A200QQX5_9MAGN) | RBFA | -0.600 | 3.50E-02 | 9.409 | 8.588 | 8.367 | 6.472 | 6.450 | 4.658 |
| Solyc12g100160.3 | 50S ribosomal protein L6 (AHRD V3.3 *** A0A2G2YBF1_CAPAN) | RP-L6 | -0.511 | 3.54E-02 | 30.872 | 27.922 | 36.867 | 20.679 | 23.878 | 22.844 |
| Solyc12g094430.1 | Glutathione S-transferase (AHRD V3.3 *** A0A200QZS2_9MAGN) | GST | -0.599 | 3.74E-02 | 51.740 | 67.059 | 61.255 | 37.116 | 36.622 | 45.514 |
| Solyc09g064500.3 | Photosystem II reaction center Psb28 protein (AHRD V3.3 *** A0A1U8EF03_CAPAN) | PSB28 | -1.254 | 3.78E-02 | 6.658 | 11.791 | 7.539 | 2.434 | 2.838 | 5.556 |
| Solyc12g009250.3 | 10 kDa chaperonin (AHRD V3.3 *** A0A1U8HGQ0_CAPAN) | GROES | -0.630 | 3.80E-02 | 130.899 | 100.459 | 170.299 | 96.153 | 91.566 | 73.696 |
| Solyc04g009030.3 | Glyceraldehyde-3-phosphate dehydrogenase (AHRD V3.3 *** A0A0A8IBT8_NICBE) | GAPA | -0.647 | 3.82E-02 | 21.485 | 13.666 | 22.072 | 10.678 | 13.494 | 12.431 |
| Solyc09g092450.3 | Long-chain acyl-CoA synthetase (AHRD V3.3 *** A0A1Z5KCI0_FISSO) | ACSL | -0.750 | 3.85E-02 | 4.025 | 3.905 | 3.958 | 3.068 | 2.560 | 1.552 |
| Solyc10g085550.3 | Enolase (AHRD V3.3 *** A0A200Q2G1_9MAGN) | ENO | -0.889 | 3.87E-02 | 13.648 | 23.011 | 10.840 | 9.913 | 8.629 | 7.498 |
| Solyc04g014270.3 | ATP-dependent 6-phosphofructokinase (AHRD V3.3 *** A0A2G2XA03_CAPBA) | PFKA | -0.440 | 3.92E-02 | 12.790 | 11.112 | 11.586 | 8.587 | 9.001 | 8.688 |
| Solyc12g099810.2 | CRT (Chloroquine-resistance transporter)-like transporter (AHRD V3.3 *** A0A1Y1I3N1_KLENI) | NA | -0.527 | 3.94E-02 | 25.733 | 26.243 | 26.772 | 17.512 | 16.255 | 20.903 |
| Solyc03g097190.3 | WEB family protein, chloroplastic (AHRD V3.3 *** A0A2G2VB89_CAPBA) | NA | -0.661 | 4.13E-02 | 11.659 | 13.208 | 10.822 | 8.201 | 9.691 | 5.106 |
| Solyc05g008600.3 | ripening regulated protein (DDTFR6/A) | ALDO | -0.471 | 4.20E-02 | 222.115 | 234.140 | 240.864 | 183.772 | 169.789 | 153.687 |
| Solyc06g066440.3 | hexokinase 2 | HK | -0.592 | 4.24E-02 | 6.206 | 5.784 | 5.340 | 4.562 | 4.026 | 3.027 |
| Solyc09g090900.4 | 3-isopropylmalate dehydratase large subunit (AHRD V3.3 *** A0A2G3CJZ4_CAPCH) | LEUC | -2.309 | 4.25E-02 | 3.943 | 5.375 | 8.651 | 1.653 | 1.976 | 0.130 |
| Solyc09g090220.3 | Pentatricopeptide repeat (AHRD V3.3 *** A0A200QEC4_9MAGN) | NA | -0.761 | 4.27E-02 | 0.981 | 0.904 | 0.897 | 0.533 | 0.520 | 0.584 |

| | | | | | | | | | | |
|---|---|---|---|---|---|---|---|---|---|---|
| Solyc12g014490.3 | 65-kDa microtubule-associated protein 1-like (AHRD V3.3 *** Q9FEV7_TOBAC),Pfam:PF03999 | PRC1 | -1.577 | 4.32E-02 | 1.484 | 2.230 | 4.634 | 1.344 | 1.106 | 0.402 |
| Solyc06g065390.4 | 50S ribosomal protein L21, chloroplastic (AHRD V3.3 *** A0A1U8GUE9_CAPAN) | RP-L21 | -0.490 | 4.37E-02 | 47.372 | 53.650 | 56.302 | 35.317 | 36.473 | 40.641 |
| Solyc10g006030.3 | 30S ribosomal protein S10, chloroplastic (AHRD V3.3 *** A0A2G2VQX8_CAPBA) | RP-S10 | -0.508 | 4.44E-02 | 27.638 | 28.737 | 34.474 | 20.958 | 20.337 | 22.771 |
| Solyc02g091580.4 | Oligopeptidase A (AHRD V3.3 *** A0A2U1NZK4_ARTAN) | PRLC | -0.412 | 4.53E-02 | 47.433 | 44.219 | 45.125 | 36.682 | 35.473 | 31.421 |
| Solyc11g011250.3 | dehydroascorbate reductase 2 | DHAR | -0.456 | 4.55E-02 | 48.084 | 44.999 | 53.363 | 34.657 | 34.280 | 38.096 |
| Solyc08g081200.3 | Short-chain dehydrogenase TIC 32, chloroplastic (AHRD V3.3 *** A0A2G2XJP0_CAPBA) | NA | -0.481 | 4.56E-02 | 18.071 | 17.153 | 20.631 | 13.871 | 15.450 | 11.121 |
| Solyc05g047550.4 | Protein kinase domain (AHRD V3.3 *** A0A200QZ79_9MAGN) | NA | -0.851 | 4.70E-02 | 1.414 | 1.389 | 2.373 | 0.842 | 1.069 | 0.972 |
| Solyc11g020060.2 | D-3-phosphoglycerate dehydrogenase (AHRD V3.3 *** A0A1J3DFK2_NOCCA) | NA | -0.499 | 4.72E-02 | 25.499 | 24.389 | 23.708 | 15.295 | 17.325 | 19.534 |
| Solyc01g098380.4 | 4-hydroxy-tetrahydrodipicolinate reductase 1, chloroplastic-like (AHRD V3.3 *** A0A1U8FAN3_CAPAN) | DAPB | -0.403 | 4.86E-02 | 31.348 | 29.347 | 30.533 | 25.069 | 22.518 | 21.819 |
| Solyc02g021000.3 | protein COFACTOR ASSEMBLY OF COMPLEX C SUBUNIT B CCB3, chloroplastic (AHRD V3.3 *** A0A2I4G7H0_9ROSI) | NA | -0.523 | 4.88E-02 | 12.348 | 11.818 | 14.621 | 9.345 | 9.714 | 8.130 |
| Solyc02g014360.3 | Pentatricopeptide repeat-containing protein (AHRD V3.3 *** A0A1U8FF53_CAPAN) | NA | -0.602 | 4.88E-02 | 3.067 | 2.408 | 2.626 | 1.968 | 1.939 | 1.471 |
| Solyc01g109160.4 | CYP74C4 | AOS | -1.099 | 4.91E-02 | 3.857 | 2.423 | 4.603 | 1.965 | 2.329 | 0.875 |
| Solyc05g055440.1 | Histone H2B (AHRD V3.3 *** A0A1U8HBK7_CAPAN) | H2B | -0.450 | 5.00E-02 | 252.947 | 222.259 | 279.774 | 177.786 | 188.751 | 188.244 |
| Solyc12g056740.2 | RNA helicase DEAD39 | NA | -0.438 | 5.00E-02 | 28.659 | 31.269 | 30.076 | 22.457 | 21.098 | 23.201 |

**Table S2. List of GO terms with top significance retrieved from the RNAseq analysis of the AM fruits.**

| | GO.ID | Term | Annotated | Significant | Expected | classic |
|---|---|---|---|---|---|---|
| O.E. Molecular function | GO:0036094 | small molecule binding | 3924 | 259 | 165.82 | 1.1E-14 |
| | GO:0043168 | anion binding | 3089 | 215 | 130.53 | 2.3E-14 |
| | GO:0016491 | oxidoreductase activity | 2396 | 176 | 101.25 | 9.6E-14 |
| | GO:0003824 | catalytic activity | 13097 | 651 | 553.44 | 2.4E-11 |
| | GO:0016829 | lyase activity | 701 | 68 | 29.62 | 1.6E-10 |
| | GO:0043177 | organic acid binding | 154 | 27 | 6.51 | 3.1E-10 |
| | GO:0043167 | ion binding | 6727 | 371 | 284.26 | 5.6E-10 |
| | GO:0050661 | NADP binding | 157 | 26 | 6.63 | 2.3E-09 |
| | GO:0016853 | isomerase activity | 346 | 41 | 14.62 | 2.8E-09 |
| | GO:0016840 | carbon-nitrogen lyase activity | 42 | 13 | 1.77 | 0.00000001 |
| U.E. Molecular function | GO:0005509 | calcium ion binding | 343 | 23 | 7.55 | 0.0000025 |
| | GO:0005515 | protein binding | 13199 | 333 | 290.52 | 0.000049 |
| | GO:0000976 | transcription cis-regulatory region bind... | 764 | 33 | 16.82 | 0.0002 |
| | GO:0001067 | transcription regulatory region nucleic ... | 769 | 33 | 16.93 | 0.00022 |
| | GO:0043565 | sequence-specific DNA binding | 1767 | 60 | 38.89 | 0.00053 |
| | GO:0031418 | L-ascorbic acid binding | 34 | 5 | 0.75 | 0.00083 |
| | GO:1990837 | sequence-specific double-stranded DNA bi... | 840 | 33 | 18.49 | 0.00103 |
| | GO:0043394 | proteoglycan binding | 10 | 3 | 0.22 | 0.00113 |
| | GO:0042393 | histone binding | 275 | 15 | 6.05 | 0.00122 |
| | GO:0030246 | carbohydrate binding | 247 | 14 | 5.44 | 0.00122 |

| | GO.ID | Term | Annotated | Significant | Expected | classic |
|---|---|---|---|---|---|---|
| O.E. Cellular component | GO:0009570 | chloroplast stroma | 1202 | 133 | 52.01 | 6.8E-24 |
| | GO:0009532 | plastid stroma | 1240 | 133 | 53.65 | 1.2E-22 |
| | GO:0009507 | chloroplast | 4138 | 299 | 179.04 | 2E-21 |
| | GO:0009536 | plastid | 4620 | 317 | 199.89 | 2.5E-19 |
| | GO:0005829 | cytosol | 5948 | 374 | 257.35 | 7.1E-17 |
| | GO:0071162 | CMG complex | 7 | 7 | 0.3 | 2.8E-10 |
| | GO:0005737 | cytoplasm | 17679 | 834 | 764.91 | 0.000000039 |
| | GO:0009941 | chloroplast envelope | 1151 | 89 | 49.8 | 0.000000075 |
| | GO:0042555 | MCM complex | 11 | 7 | 0.48 | 0.000000079 |
| | GO:0009526 | plastid envelope | 1648 | 116 | 71.3 | 0.00000012 |
| U.E. Cellular component | GO:0012506 | vesicle membrane | 849 | 38 | 19.41 | 0.00007 |
| | GO:0030659 | cytoplasmic vesicle membrane | 808 | 36 | 18.47 | 0.00012 |
| | GO:0016514 | SWI/SNF complex | 26 | 5 | 0.59 | 0.00027 |
| | GO:0098936 | intrinsic component of postsynaptic memb... | 17 | 4 | 0.39 | 0.00051 |
| | GO:0099055 | integral component of postsynaptic membr... | 8 | 3 | 0.18 | 0.00061 |
| | GO:0099503 | secretory vesicle | 475 | 23 | 10.86 | 0.00066 |
| | GO:0000785 | chromatin | 488 | 23 | 11.16 | 0.00094 |
| | GO:0099240 | intrinsic component of synaptic membrane | 21 | 4 | 0.48 | 0.00119 |
| | GO:0001669 | acrosomal vesicle | 36 | 5 | 0.82 | 0.00129 |
| | GO:0005886 | plasma membrane | 6238 | 173 | 142.6 | 0.0018 |

| | GO.ID | Term | Annotated | Significant | Expected | classic |
|---|---|---|---|---|---|---|
| O.E. Biological processes | GO:0044281 | small molecule metabolic process | 4042 | 313 | 170.25 | 7.1E-30 |
| | GO:0019752 | carboxylic acid metabolic process | 2386 | 216 | 100.5 | 2E-28 |
| | GO:0043436 | oxoacid metabolic process | 2857 | 236 | 120.34 | 1.8E-25 |
| | GO:0006082 | organic acid metabolic process | 3051 | 245 | 128.51 | 8.9E-25 |
| | GO:0044283 | small molecule biosynthetic process | 1783 | 158 | 75.1 | 1.3E-19 |
| | GO:0006520 | cellular amino acid metabolic process | 805 | 93 | 33.91 | 7.8E-19 |
| | GO:0055086 | nucleobase-containing small molecule met... | 761 | 86 | 32.05 | 6.7E-17 |
| | GO:0006753 | nucleoside phosphate metabolic process | 626 | 76 | 26.37 | 9.3E-17 |
| | GO:0032787 | monocarboxylic acid metabolic process | 1560 | 137 | 65.71 | 1.1E-16 |
| | GO:0009117 | nucleotide metabolic process | 621 | 75 | 26.16 | 2E-16 |
| U.E. Biological processes | GO:0050896 | response to stimulus | 15114 | 391 | 338.67 | 0.00000068 |
| | GO:0065007 | biological regulation | 13786 | 363 | 308.91 | 0.00000068 |
| | GO:0006873 | cellular ion homeostasis | 574 | 32 | 12.86 | 0.0000026 |
| | GO:0030003 | cellular cation homeostasis | 530 | 30 | 11.88 | 0.0000039 |
| | GO:0055082 | cellular chemical homeostasis | 721 | 36 | 16.16 | 0.0000074 |
| | GO:0010035 | response to inorganic substance | 3727 | 120 | 83.51 | 0.000017 |
| | GO:0006875 | cellular metal ion homeostasis | 409 | 24 | 9.16 | 0.00002 |
| | GO:0050789 | regulation of biological process | 12747 | 332 | 285.63 | 0.000025 |
| | GO:0055080 | cation homeostasis | 837 | 37 | 18.76 | 0.000075 |
| | GO:0006911 | phagocytosis, engulfment | 36 | 6 | 0.81 | 0.00014 |

**Table S3. Top 25 genes overexpressed in the AM fruits contributing to the chloroplast and plastid-related GO terms of the cellular component.**

| gene | description | logFC | FDR |
|---|---|---|---|
| Solyc09g059020.4 | Quinone-oxidoreductase QR1, chloroplastic (AHRD V3.3 *** QR1_TRIVS) | -9.275 | 7.00E-197 |
| Solyc03g044330.1 | Acetolactate synthase (AHRD V3.3 *** C0L093_TOBAC) | -9.673 | 1.89E-92 |
| Solyc09g059060.1 | Quinone-oxidoreductase QR1, chloroplastic (AHRD V3.3 *-* QR1_TRIVS),Pfam:PF13602 | -12.768 | 4.29E-73 |
| Solyc05g050980.3 | 3-phosphoshikimate 1-carboxyvinyltransferase (AHRD V3.3 *** A0A0M8KSM3_NICAT) | -4.822 | 2.05E-70 |
| Solyc11g066890.1 | Arogenate dehydratase (AHRD V3.3 *** A0A2G3BBC7_CAPCH) | -3.002 | 4.00E-49 |
| Solyc09g059070.3 | Quinone-oxidoreductase chloroplastic-like (AHRD V3.3 *-* A0A2K3P901_TRIPR) | -11.991 | 3.95E-48 |
| Solyc06g075010.4 | 60 kDa chaperonin (AHRD V3.3 *** B2IXD2_NOSP7) | -5.960 | 9.48E-47 |
| Solyc10g084400.2 | Glutathione S-transferase (AHRD V3.3 *** Q76KW1_PEA) | -2.941 | 9.92E-43 |
| Solyc08g067310.1 | Non-specific serine/threonine protein kinase (AHRD V3.3 *** A0A2G2W7U9_CAPBA) | -4.071 | 2.74E-38 |
| Solyc04g011390.1 | Histone H4 (AHRD V3.3 *-* F2E7L1_HORVV) | -4.686 | 2.17E-37 |
| Solyc01g008920.4 | NAD(P)-binding Rossmann-fold superfamily protein (AHRD V3.3 *** A0A2U1L7S8_ARTAN) | -2.977 | 1.35E-30 |
| Solyc06g074790.2 | Histone H2B (AHRD V3.3 *** A0A2G3CAQ8_CAPCH) | -3.575 | 5.87E-26 |
| Solyc10g007100.3 | Protein DETOXIFICATION (AHRD V3.3 *** A0A218WWW8_PUNGR) | -3.341 | 1.20E-25 |
| Solyc03g114500.4 | Enolase (AHRD V3.3 *** A0A2G2WR88_CAPBA) | -5.816 | 2.97E-25 |
| Solyc03g025840.3 | Cytochrome b561/ferric reductase transmembrane protein family (AHRD V3.3 *** A0A178UZ09_ARATH) | -2.894 | 1.15E-23 |
| Solyc06g083440.3 | Cytochrome b5 (AHRD V3.3 *** A0A2G3CCB6_CAPCH) | -2.895 | 2.07E-22 |
| Solyc06g073190.3 | fructokinase 2 | -1.687 | 9.22E-22 |
| Solyc10g083360.2 | Calmodulin-binding family protein, putative, expressed (AHRD V3.3 *** Q2QXN6_ORYSJ) | -1.778 | 1.06E-20 |
| Solyc06g005160.4 | cytosolic ascorbate peroxidase 1 | -1.731 | 5.71E-19 |
| Solyc06g005430.1 | Histone H4 (AHRD V3.3 *-* F2E7L1_HORVV) | -1.903 | 8.15E-19 |
| Solyc10g005300.3 | Serine/threonine-protein kinase PBS1 (AHRD V3.3 *** A0A2G2VSF8_CAPBA) | -3.347 | 1.55E-18 |
| Solyc05g053100.3 | Dihydrolipoyl dehydrogenase-like protein (AHRD V3.3 *** A0A2K3PDU3_TRIPR) | -2.126 | 6.86E-18 |
| Solyc09g065180.3 | binding protein precursor AF106660 | -2.061 | 3.65E-17 |
| Solyc11g056680.1 | Leucine-rich repeat receptor-like protein (AHRD V3.3 *** H6V788_MALDO) | -2.784 | 6.79E-17 |
| Solyc04g049350.4 | chorismate synthase 1 precursor | -1.543 | 1.34E-16 |

**Table S4. List of primers used in this work.**

| Name | Sequence | Use |
|---|---|---|
| M13MAY01ALS1aOF | GCGCCGTCTCACTCGAATGGCGGCTGCTGCCTCACC | Cloning |
| M13MAY02ALS1aOR | GCGCCGTCTCATGACACTTGACCTGTAATAGCAACAATCGG | Cloning |
| M13MAY03ALS1bOF | GCGCCGTCTCAGTCAAGGAGGATGATTGGTAC | Cloning |
| M13MAY04ALS1bOR | GCGCCGTCTCAGCCTCAGCTCCTCACTTGATTG | Cloning |
| M13MAY05ALS2OF | GCGCCGTCTCAAGGCGATTTGTGGAGCTTACAGG | Cloning |
| M13MAY06ALS2OR | GCGCCGTCTCACCTCCCAACAGCCGCACCTAT | Cloning |
| M13MAY07ALS3OF | GCGCCGTCTCAGAGGCCGGGTGAGATTGTGG | Cloning/Genotyping |
| M13MAY08ALS3OR | GCGCCGTCTCACTCGAAGCTCAATAGGAACATCTCCCGTCGCC | Cloning |
| M13OCT01_PE8F1 | GCGCCGTCTCACTCGGGAGTCCCTAATGATATTGTTCATG | Cloning |
| M13OCT02_PE8R1 | GCGCCGTCTCACTCGCATTCTTCTTTTGCACTGTGAATGATTAG | Cloning |
| M12MAY03TermE8F1 | GCGCCGTCTCGCTCGGCTTGAATAAGAATAATAATG | Cloning |
| M12MAY04TermE8R1 | GCGCCGTCTCGCTCGAGCGCGTAAATTAGATAAGGAAAAC | Cloning |
| M12ENE24TmtbR1 | GCGCCGTCTCGCTCGAGCGTCGCAAAAACTATATGCTCTC | Genotyping |
| MV18OCT58 qPCR SlAct7 F | CCTCAGCACATTCCAGCAG | RT-qPCR actin |
| MV18OCT59 qPCR SlAct7 R | CCACCAAACTTCTCCATCCC | RT-qPCR actin |
| MV18OCT60 qPCR 3UTRMtb F | cagccatagaaggctaacacc | RT-qPCR cALS |
| MV18OCT61 qPCR SlmutALS R | acactcctgggccatacttg | RT-qPCR cALS |
| MV18OCT62 qPCR SlMyb12 F | tgaagaagcaacaacaatgga | RT-qPCR cMYB |
| MV18OCT63 qPCR 3UTRE8 R | ctcaaacatttgcttcaaattca | RT-qPCR cMYB |
| MV19APR01 qPCR SlMyb12end F2 | catagtaccattgtatcttggcttt | RT-qPCR eMYB |
| MV18OCT65 qPCR SlMyb12end R | tcccacattccaaatataaaacg | RT-qPCR eMYB |
| MV18OCT66 qPCR SlALSend F | gcgatgggagatgttcctatt | RT-qPCR eALS |
| MV18OCT67 qPCR SlALSend R | acaacagccacaacaagcaa | RT-qPCR eALS |
